# Functional Contribution of Cancer-Associated Fibroblasts in Glioblastoma

**DOI:** 10.1101/2022.04.07.487495

**Authors:** Phillip M. Galbo, Yang Liu, Mou Peng, Yao Wei, Anne Tranberg Madsen, Sarah Graff, Cristina Montagna, Jeffrey E. Segall, Simone Sidoli, Xingxing Zang, Deyou Zheng

## Abstract

The abundance and biological contribution of cancer associated fibroblasts (CAFs) in glioblastoma are poorly understood. Here, we applied single-cell RNA sequencing and spatial transcriptomics analyses to identify and characterize CAFs in human glioblastoma tumors and then performed functional enrichment analysis and *in vitro* assays to investigate their interactions with malignant glioblastoma cells. We found that CAF abundance was significantly correlated with tumor grade, poor clinical outcome, and activation of extracellular matrix remodeling, using three large databases containing bulk RNA-sequencing data and clinical information. Proteomic analysis of the CAFs and their secretome revealed fibronectin (FN1) as a strong candidate mediating CAF functions. This was validated using *in vitro* cellular models, which demonstrated that CAF conditioned media and recombinant FN1 could facilitate the migration and invasion of glioblastoma cells. In addition, we showed that CAFs were more abundant in the mesenchymal-like state (or subtype) than in other states of glioblastomas, while cell lines resembling the proneural-state responded to the CAF signaling better in terms of the migratory and invasive phenotypes. Investigating the *in-situ* expression of gene markers specifically associated with CAFs and mesenchymal malignant cells further indicated that CAFs were enriched in the perinecrotic and pseudopalisading zones of human tumors, where mesenchymal-like glioblastoma cells co-resided and thus likely interacted. Overall, this study characterized the molecular features and functional impacts of CAFs in glioblastoma, alluding to a novel cell-to-cell interaction axis mediated by CAFs in the glioblastoma microenvironment.

## Introduction

About one-third of all brain tumors are gliomas, while glioblastoma multiforme (GBM) is the most common malignant brain tumor diagnosed in adults (*1*). With very limited therapeutic options, there is a dire need for better understanding of the underlying mechanisms governing GBM pathogenesis and treatment responses (*2*). The tumor microenvironment (TME) is a major barrier in the effective management of patients diagnosed with glioblastoma. Specifically, studies have demonstrated that the TME confers resistance to targeted therapies (*3*), intra-tumoral heterogeneity (*4*), and malignant-cell invasion (*5-7*). Thus, studying glioblastoma-to-TME interactions can help delineate novel mechanisms associated with pathogenesis.

In epithelial cancers, malignant cell interactions with CAFs can confer a wide range of biological processes that contribute to disease progression. These include tumorigenesis (*8, 9*), angiogenesis (*10*), therapeutic resistance (*11*), anti-tumoral immunity (*12*), inflammation (*13*), and malignant-cell invasion (*14-16*). The molecular characteristics and functional roles of CAFs in glioblastoma, however, remain to be established. It is conceivable that a better understanding of the interaction between CAFs and malignant cells in glioblastoma could elucidate novel mechanisms mediating tumorigenesis and progression, uncovering potentially new therapeutic strategies.

Towards this goal, we have performed advanced bioinformatic analysis of single cell RNA-sequencing (scRNA-seq) (*17*) and spatial transcriptomics data from human glioblastoma tumors to identify and characterize the molecular features of CAFs. Using signature genes of the CAFs and Cancer Genome Atlas (*18, 19*) datasets, we further carried out multiple-level orthogonal analyses to study how CAF enrichment scores were associated with tumor grade, patient survival, GBM state heterogeneity, and abnormal cellular processes such as extracellular matrix remodeling. Proteomic analysis and *in vitro* cell modeling identified fibronectin 1 (FN1) as a top soluble factor of the CAF-secretome that could enhance migration and invasion of malignant cells. Lastly, RNA-seq and *in situ* hybridization analysis indicated that CAFs were enriched in the perinecrotic/pseudopalisading zones of human tumors, adjacent to mesenchymal-like glioblastoma cells. Collectively, our work describes a novel functional interaction in the glioblastoma TME involving CAFs that may participate in several pathologies.

## Results

### Identification of CAFs as a novel cell type important to disease progression in glioma

Stromal cells in TME can exert a tumor-promoting or -suppressive effect. Thus, we first sought to evaluate how the abundance of different TME cell types is correlated with clinical outcome in glioma. We used the program EPIC (Estimating the Proportions of Immune and Cancer Cells) (*20*) to deconvolute cell type compositions in three RNA-seq datasets obtained from bulk tumors of either low grade gliomas (LGG) or glioblastoma that are available in The Cancer Genome Atlas (TCGA) and Chinese Glioma Genome Atlas (CGGA) databases (*18, 19*). Applying univariate cox proportional hazard analysis to the estimated cell type proportions in the TCGA data, we found that CAF abundances were associated with poor prognosis in both LGG and glioblastoma tumors (**Figure S1a, b**). The findings were reproduced when CGGA 325 and CGGA 693 were analyzed (**Figure S1a, b**). In addition, Kaplan-Meier estimator analysis suggests that patients diagnosed with either LGG or glioblastoma with a high proportion of CAFs had a significantly shorter survival, compared to patients with a low proportion of CAFs in their tumors (**Figure S1c-h**). Collectively, these analyses implicated CAFs as a novel cell type that might contribute to a more aggressive disease state in both LGG and GBM.

### Identification and molecular characterization of CAFs in glioblastoma

We next searched for additional and more direct evidence in support of our findings because the existence of CAFs in glioblastoma is not well established, partially due to the lack of unique markers. Single cell RNA-sequencing (scRNA-seq) datasets provide a good means to address this, but standard data analysis can miss rare cell types. This is important because the EPIC results suggest that CAFs are <5% in the bulk tumors. Thus, we used RaceID3, a software designed specifically to identify outlier (i.e., rare) cells in scRNA-seq datasets (*21*), to re-analyze previously published human glioblastoma scRNA-seq datasets using 10,000 randomly selected non-malignant cells (*17*). The analysis identified a total of 548 outlier cells, among which was a cluster of 187 cells closely resembling the transcriptomic characteristics of CAFs described in other types of solid tumors (*22*). Specifically, these cells highly expressed CAF activated markers *ACTA2* (*α-SMA*), *VIM, LOX*, and *CAV1* (**Figure 1a**) (*38*). They also expressed highly an array of collagens commonly produced by CAFs, including *COL1A1, COL4A1, COL5A1*, and *COL6A1* (**Figure 1a**) (*22*). Furthermore, they showed minimal or no expression of pan-immune cell maker *CD45*, macrophage markers *CD14* and *CSF1R*, B cell markers *CD79A/B*, endothelial markers *PECAM1* (*CD31*) and *VWF*, pericyte marker *RGS5*, and astrocytic marker *GFAP* (**Figure 1a**). Comparison of their top marker genes to 1,368 scRNA-seq datasets in the PanglaoDB (*23*) further supported that these cells were most similar to fibroblast cells (**Figure 1b**). Based on these data, we considered those 187 outlier cells as CAFs (or CAF-like cells).

**Figure 1.**
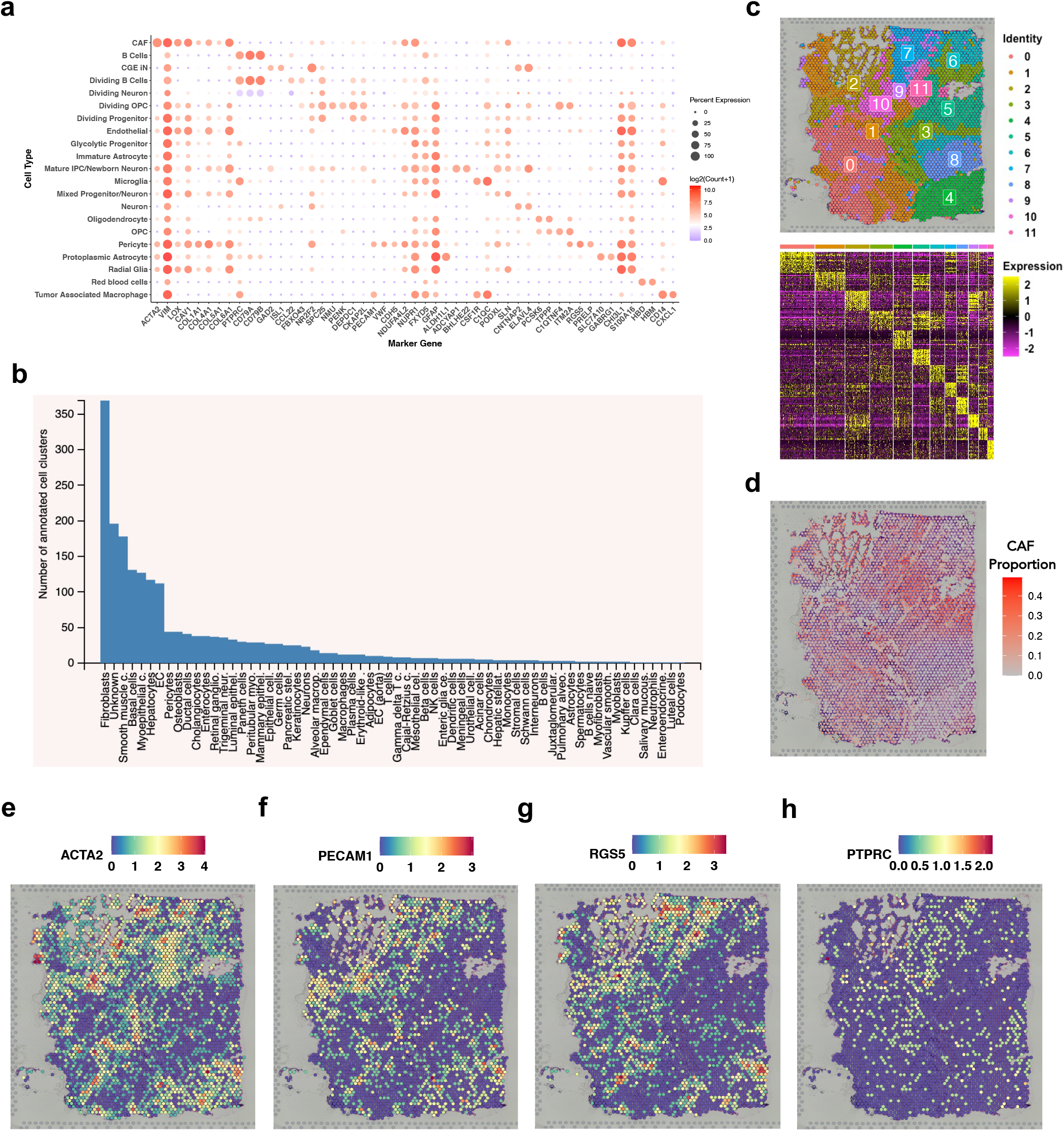
Identification and molecular characterization of CAFs in glioblastoma. (**a**) Bubble-plot indicating the average expression of selected marker genes for the cell-types identified in human glioblastomas. The size of the circle corresponds to the percent of cells that express a marker while the color indicates the log_2_(count+1). (**b**) Bar plot indicating the number of cell clusters in the PanglaoDB with markers significantly overlapping with the CAF markers identified here. (**c**) Spatial Dimension Plot (dimplot) of a human glioblastoma tissue sample indicating distinct transcriptomic regions (top) and heatmap displaying the top 20 marker genes (bottom). (**d**) Density heatmap showing the estimated CAF proportions throughout the glioblastoma tissue. (**e-h**) Spatial dimplot showing gene expression patterns for (**e**) *ACTA2*, (**f**) *PECAM1 (CD31*), (**g**) *RGS5*, and (**h**) *PTPRC (CD45*).

To further characterize the features associated with the CAFs, we set out to identify and profile CAF-enriched niches using spatial transcriptomics data. Our analysis identified 12 transcriptionally distinct regions in a human glioblastoma sample (**Figure 1c**). We then applied MuSiC, a method that leverages cell-type specific gene expression from scRNA-seq data to deconvolute cell type estimates from bulk RNA-seq data (*24*), to identify which of the 12 regions was enriched for CAFs. The result revealed that CAFs were present in several regions, but the enrichment was high in the regions 5 and 11 (**Figure 1d**), supported by high expression of the CAF-specific marker genes from the scRNA-seq analysis above. Specifically, regions 5 and 11 showed high expression of *ACTA2* (*α-SMA*), but reduced expression of *PECAM1* (*CD31*), *RGS5*, and *PTPRC* (*CD45*) (**Figure 1e-h**). We also analyzed the expression of genes previously found to strongly associated with vascular proliferation (*25*) and found they showed a lack of expression in the regions 5 and 11 (**Figure S2a**), indicating that the two regions likely did not contain a significant number of vascular cells involved in angiogenesis. The regions also highly expressed other markers associated with CAFs, identified in our scRNA-seq analysis, including *GJB2, PTX3, SERPINE1, ACTG2, AKAP12*, and *ITGA5* (**Figure S2 b-g**).

Collectively, results above support strongly that CAFs are sufficiently distinct from vascular and other cell types in glioblastoma, suggesting a need to further study their unique biological functions in mediating GBM pathogenesis.

### CAFs association with tumor grade and clinical in glioma

Previous studies suggest that CAFs can either promote (*11-13, 16*) or suppress (*26-28*) the survival of malignant cells, depending on the tumor types. We thus sought to evaluate if CAFs are tumor-promotive or -suppressive, by first assessing if CAFs correlate with tumor grade. This was accomplished by curating a CAF-specific gene signature from the scRNA-seq analysis to calculate a CAF-enrichment score for each human bulk RNA-seq sample in the TCGA and CGGA cohorts. This analysis revelated that CAF scores were significantly higher in grade IV compared to grade II and III gliomas in all three datasets (**Figure 2a-c**). To further address if CAFs are tumor-promotive or -suppressive, we evaluated the prognostic utility of the CAF-specific gene signature from our GBM scRNA-seq analysis. This approach is an improvement over the EPIC analysis described above because the CAF gene signature in EPIC was derived from melanoma tumors and the gene expression profile of glioblastoma-CAFs showed some differences from that of melanoma-CAFs, likely reflecting distinct TMEs. We compared the prognostic utility of the CAF gene signature identified here by us and 20 gene signatures for other cell types identified previously by Bhaduri *et al*. in the same glioblastoma samples (*17*). Specifically, an enrichment score was calculated for each of the 21 cell type gene signatures in the TCGA or CGGA samples. The enrichment scores were then related to patient outcomes with a univariate cox-proportional hazard model. Consistent with literature (*29*), we observed that tumor associated macrophages and microglia were significantly correlated with a poor prognosis in all three datasets from either LGG or glioblastoma tumors (**Figure 2d, e**), indicating the robustness of our analysis. The results also indicate that CAF scores were significantly correlate with a poor clinical outcome in all datasets (**Figure 2d, e**). Furthermore, in a multivariate analysis, the impact of CAF enrichment score on survival of glioma patients was found to be independent of age, gender, isocitrate dehydrogenase (IDH1) mutation status, and grade (covariates know to be associated with clinical outcome in glioma) (**Table S1**).

**Figure 2.**
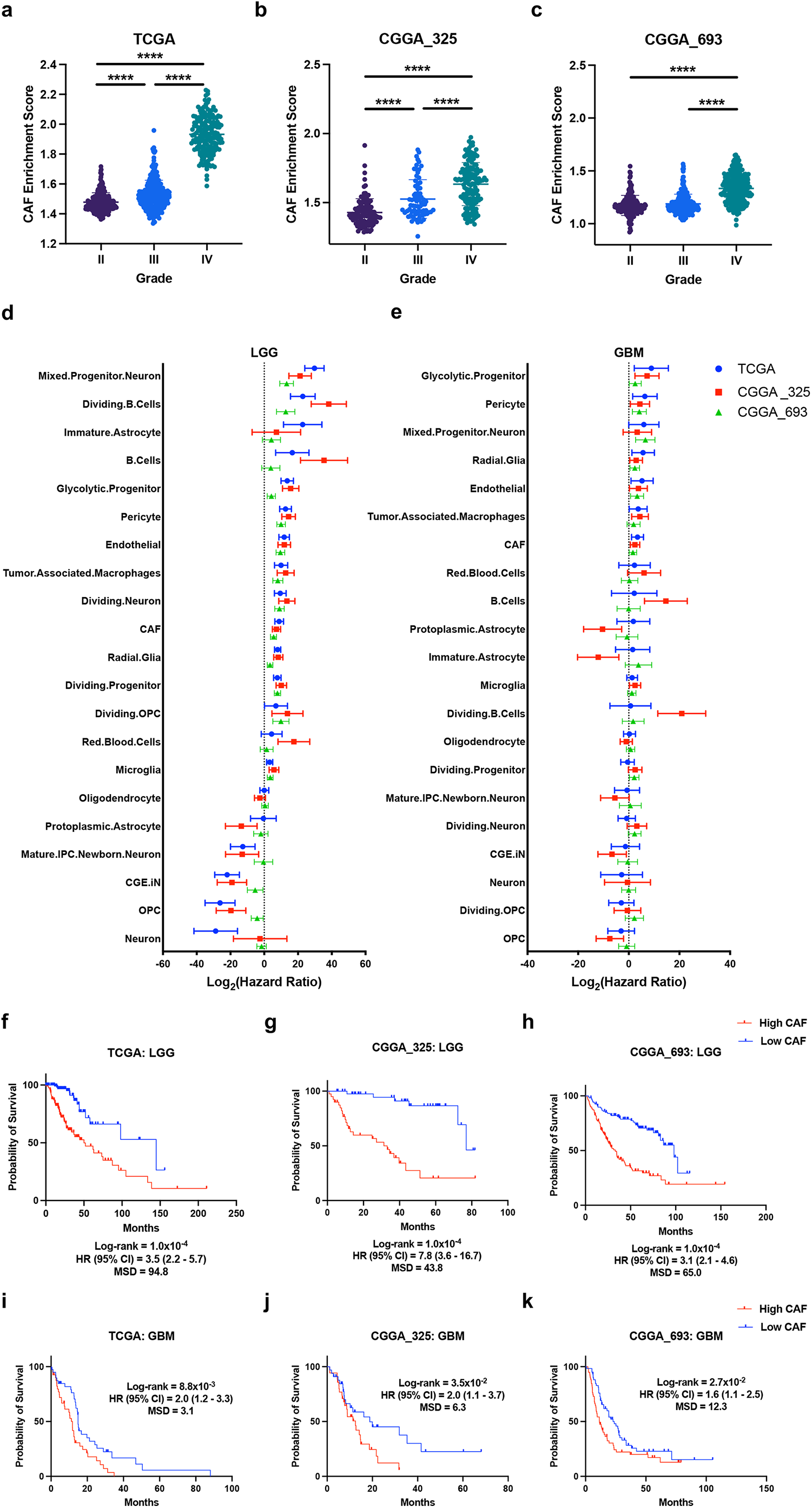
CAF association with tumor grade and clinical outcome in glioma. (**a-c**) Association between tumor grade and CAF enrichment score in (**a**) TCGA (Grade II, n=248; Grade III, n=261; Grade IV n=173), (**b**) CGGA 325 (Grade II, n=109; Grade III, n=72; Grade IV, n=144), and (**c**) CGGA 693 (Grade II, n=188; Grade III, n=255; Grade IV n=249) datasets (data displayed as mean ± S.D; One-way ANOVA: **** < 0.0001). (**d-e**) Hazard ratios associated with cell-type-specific enrichment scores in three datasets describing (**d**) LGG or (**e**) glioblastoma samples (data displayed as median with range). (**f-h**) Kaplan-Meier plots indicating percent survival over time (months) among LGG patients with either a high- or low-CAF enrichment score in (**f**) TCGA (High CAF; n=127; Low CAF, n=127), (**g**) CGGA 325 (High CAF, n=45; Low CAF, n=45), and (**h**) CGGA 693 (High CAF, n=111; Low CAF, n=111) datasets (MSD, median survival difference; HR, hazard ratio; CI, confidence interval). (**i-k**) Kaplan-Meier plots indicating percent survival over time (months) among glioblastoma patients with either a high- or low-CAF enrichment score in (**i**) TCGA (High CAF, n=40; Low CAF, n=40), (**j**) CGGA 325 (High CAF, n=36; Low CAF, n=36), and (**k**) CGGA 693 (High CAF, n=62; Low CAF, n=62) datasets (MSD, median survival difference; HR, hazard ratio; CI, confidence interval).

We further assessed the prognostic utility of the CAF-specific gene signature by Kaplan-Meier Estimator analysis. In all three independent datasets, we found that LGG patients with a high CAF score had a significantly shorter overall survival compared to those with a low score (**Figure 2f-h**). In a multivariate analysis, the impact of CAF score on survival in LGG was found to be independent of age, gender, isocitrate dehydrogenase (IDH1) mutation status, and neoplasm grade (covariates known to be associated with clinical outcome in LGG) (**Table S2**). Among patients diagnosed with glioblastoma, across three datasets, Kaplan-Meier Estimator analysis showed that patients with a high CAF score also had a shorter overall survival than patients with a low score (**Figure 2i-k**). However, in multivariate analysis, the impact of CAFs was only moderately associated with survival in glioblastoma after adjusting for age, gender, IDH mutation status, and O-G-methyl guanine-DNA methyl transferase (MGMT) methylation (covariates known to be associated with clinical outcome in glioblastoma) (**Table S3**). In addition, we observed that patients with a high CAF score in the TCGA cohort were more susceptible to a shorter disease-free progression than patients with a low CAF score (**Figure S3a**). However, CAF scores did not show a significant difference between primary- and reoccurring-glioblastoma cases (**Figure S3b-d**).

Collectively, these results demonstrate the prognostic utility of our CAF gene signature in both LGG and glioblastoma and suggest that CAFs may promote malignant cell survival in glioblastoma.

### CAFs contribute to malignant cell migration and invasion in glioblastoma

Having shown that CAFs are significantly associated with disease progression, we turned to the potential biological mechanisms, by identifying molecular and cellular pathways that are differentially active between the glioblastomas with high and low CAF scores. A principal component analysis (PCA) of the transcriptomes of the TCGA and CCGA samples revealed that glioblastomas with a high CAF score were closely associated but separated from glioblastomas with a low CAF score (**Figure 3a-c**), suggesting a global gene expression difference among the two groups. Next, we performed gene set enrichment analysis (GSEA) (*30*) to identify gene sets that were enriched (i.e., more active) in one of the two groups, followed by enrichmentmap analysis (*31*) to group gene sets to common biological themes. The analysis uncovered key biological processes with significantly higher activities in glioblastomas with a high CAF score, such as inflammation (i.e., Inflammatory Response and Signaling via Interleukins), extracellular matrix (ECM) remodeling, and metabolism (i.e., Metabolic Activity) (**Figure 3d**). Upon a closer examination of the individual gene sets under the ECM remodeling, we observed several gene sets related to cell migration and invasion (i.e., Anatassiou Multicancer Invasiveness Signature, WU Cell Migration, and Wang Tumor Invasiveness Up) (**Figure 3d**), consistent with existing literature linking ECM to enhanced tumor invasion (*32*). Interestingly, all the identified biological themes among GBMs with a high CAF score are strongly associated with a more aggressive disease state (*33*). Conversely, glioblastomas with a low CAF score showed greater activities in gene sets related to cellular cycle regulation (i.e., Cell Cycle, DNA Repair, and Telomere Maintenance), synaptic activity, and RNA processing (**Figure 3d**). These results indicate that CAFs reside in a glioblastoma microenvironment that favors inflammation, cell migration, and cell invasion, while glioblastomas lacking CAFs are strongly associated with cell proliferation.

**Figure 3.**
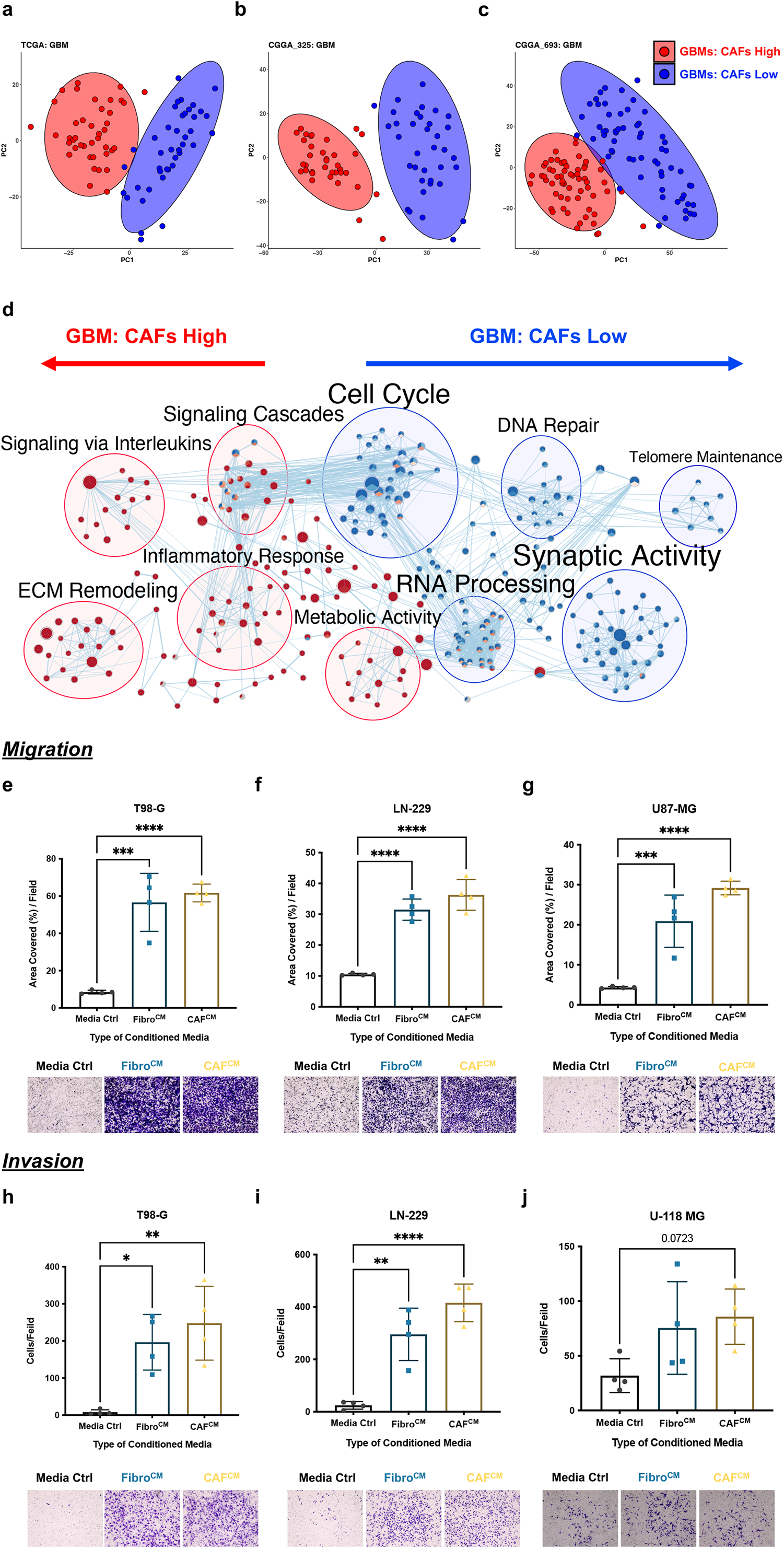
CAFs contribute to malignant cell migration and invasion in glioblastoma. (**a-c**) PCA plots of RNA-seq samples from glioblastomas with high and low CAF enrichment scores for (**a**) TCGA (High CAF, n=40; Low CAF, n=40), (b) CGGA 325 (High CAF, n=36; Low CAF, n=36), and (**c**) CGGA 693 (High CAF, n=62; Low CAF, n=62) datasets. Ellipse defines a region that contains 95% of all samples belonging to a particular group. (**d**) Networks of REACTOME terms enriched or depleted in glioblastomas with high CAF enrichment score compared to glioblastomas with low CAF score. Nodes are terms enriched (red circles) or depleted (blue circles) among genes expressed higher in glioblastomas with high CAF enrichment scores, while edges link terms with overlapping genes. Connected nodes with similar functions are further summarized by a more generalized biologically relevant term using Enrichment map. Each node is composed of three parts corresponding to data for TCGA, CGGA 325, and CGGA 693, and thus the color indicates how similar the enrichment was observed. (**e-g**) Transwell migration results for glioblastoma cell lines exposed to conditioned media, (**e**) T98-G, (**f**) LN-229, and (**g**) U87-MG. (**h-j**) Transwell invasion results for glioblastoma cell lines exposed to conditioned media, (**h**) T98-G, (**i**) LN-229, and (**j**) U-118 MG. Data were generated from n=4 independent experiments and displayed as mean ± S.D. One-way ANOVA: * < 0.05, ** < 0.01, *** < 0.001, and **** < 0.0001.

To study this *in vitro*, we obtained a primary CAF cell line (referred as “GBM-CAFs”) established from human glioblastoma tumors. The CAF characteristics of this cell line were first validated by the expression of several markers via immunocytochemistry, including high expression of ACTA2 (α-SMA) and VIM but lacking expression of the astrocytic marker GFAP (**Figure S4a-c**), in line with our scRNA-seq data (**Figure 1a**). We next carried out proteomic analysis of the whole cell lysates (WCL) of the GBM-CAFs, along with the IMR90 fibroblast cell line. The WCL data was analyzed comparatively with two other WCL datasets, one for CAFs and adjacent fibroblasts (AF) from human oral tongue squamous cell carcinoma (OTSCC) (*34*) and the other for resting and activated CD4+ T Cells (*35*). Using those external datasets as “positive” and “negative” controls, respectively, we observed that GBM-CAFs and OTSCC-CAFs were clustered together but away from CD4+ T cell populations, by both PCA (**Figure S4d**) and expression correlation analysis (**Figure S4e**). Taken together, these data confirm that the GBM-CAFs are indeed a CAF cell line and thus a suitable model for investigating CAF functions.

We used the GBM-CAFs to test if CAFs contribute to malignant cell migration and invasion *in vitro*. This was accomplished by harvesting conditioned media from the IMR90 fibroblast cell line (“Fibro^CM^”) and the GBM-CAF cell line (“CAF^CM^”) and using them to study effects on cell migration. Compared to control media (“Media Ctrl”), both the Fibro^CM^ and CAF^CM^ facilitated the migration of three human glioblastoma cell lines (T98-G, LN-229, and U87-MG) (**Figure 3 e-g**). However, neither Fibro^CM^ nor CAF^CM^ significantly facilitated the migration of the U-118 MG cells (data not shown). In invasion assay, we found that Fibro^CM^ and CAF^CM^ facilitated the invasion of T98-G and LN-229 (**Figure 3f-e**), but not U-118 MG (**Figure 3j**) or U87-MG (data not shown).

Collectively, the *in vitro* models indicate that CAFs contribute to malignant cell migration and invasion. Ultimately, the collection of these findings alludes to key biological functions of CAFs in glioblastoma pathogenesis that have not been previously described.

### CAF-derived fibronectin facilitates glioblastoma cell migration and invasion

Since both the Fibro^CM^ and CAF^CM^ facilitated the migration and invasion of several glioblastoma cell lines, we performed proteomic analysis to identify highly abundant secreted factors that could potentially mediate this phenotype. A proteomic screening uncovered 1,559 and 1,051 proteins in the Fibro^CM^ and CAF^CM^, respectively. To narrow down a short list of strong candidates, we designed a scoring method that took into consideration abundance ranks, relative expression in the scRNA-seq data, correlation with patient survival, and several other critical features (see Methods). As a result, a total of 44 and 37 proteins were identified uniquely in the Fibro^CM^ and CAF^CM^, respectively, with 152 proteins in common (**Figure 4a**). Since both Fibro^CM^ and CAF^CM^ induced cell migration and invasion, we decided to focus on the common ones (**Figure 4a)**. The number one top ranked protein is fibronectin (FN1) in both the Fibro^CM^ and CAF^CM^ (**Figure 4b, c**); however other top ranked proteins, including COL1A1, COL6A1, SPARC, and SERPINE1, are also known to be associated with CAF functions in other cancers (*22*). FN1 has been associated with migration and invasion in GBM (*36, 37*). Interestingly, across the TCGA and CGGA datasets, *FN1* expression was significantly and positively correlated with CAF enrichment scores (**Figure 4d-f**). This correlation was confirmed at the protein level, when we examined the proteomic data from 99 human treatment naïve glioblastomas analyzed by the Clinical Proteomic Tumor Analysis Consortium (CPTAC) (*38*) (**Figure 4g**). The spatial transcriptomic data also showed that *FN1* is expressed in the regions 5 and 11 (**Figure 4h**), where CAFs were enriched, among a few other locations (**Figure 1e**). In terms of cell type specificity, CAFs exhibited the highest expression of *FN1* and contributed the fourth most to the total *FN1* expression among all the cell types detected in the glioblastoma scRNA-seq samples (**Figure 4i**). Lastly, a ligand-receptor analysis (of the scRNA-seq dataset) suggested that CAFs could communicate with malignant cells via the FN1 signaling network in glioblastoma (**Figure 4j**). Collectively, these results suggest that CAFs contribute significantly to the FN1 signaling in GBM and is a great candidate for mediating the migration and invasion phenotype observed above.

**Figure 4.**
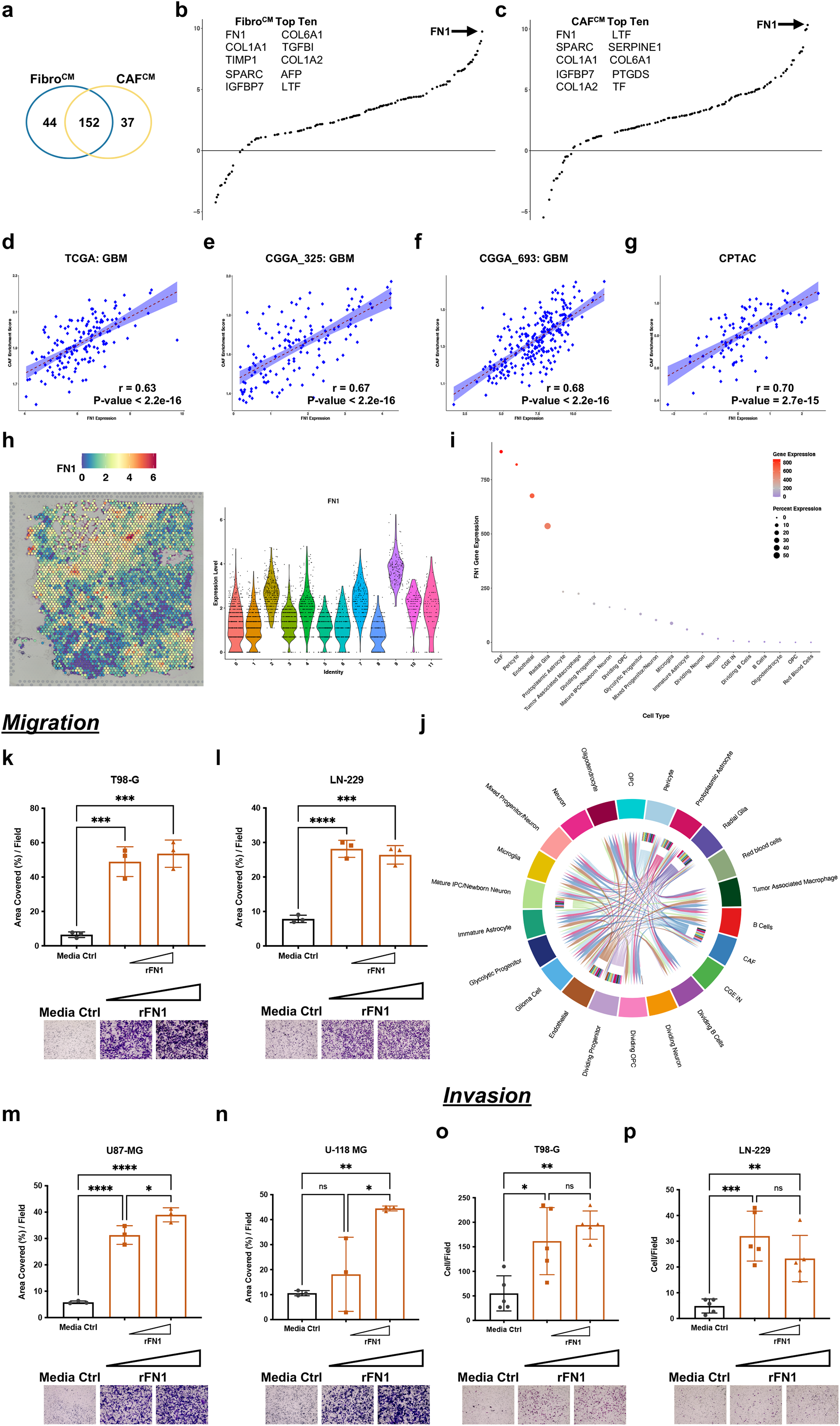
CAF-derived fibronectin facilitates glioblastoma cell migration and invasion. (**a**) Venn diagram displaying the numbers of top ranked proteins in Fibro^CM^ and CAF^CM^. (B-C) Ranks of proteins detected in (**b**) Fibro^CM^ and (**c**) CAF^CM^, with the top 10 listed. The x-axis represents the rank of proteins from low (left) to high (right), while the y-axis indicates the average protein expression across two independent replicates. (**d-f**) Pearson’s correlation between CAF enrichment scores and *FN1* expression (FPKM) in glioblastoma samples, (**d**) TCGA (n=161), (**e**) CGGA_325 (n=144), and (**f**) CGGA_693 (n=249) datasets. (**g**) Pearson’s correlation between CAF enrichment scores and FN1 protein expression in the CPTAC (n=99) dataset. (**h**) Spatial Dimension Plot displaying *FN1* expression. (**i**) Average *FN1* expression across cell-types in the scRNA-seq analysis. Size of the circle indicates individual contribution of a cell type to the total *FN1* expression, with colors indicate mean expression. (**j**) Cell-to-cell FN1 interaction network among cell types in the scRNA-seq dataset. (**k-n**) Transwell migration results for glioblastoma cell lines exposed to 10 ng/mL or 20 ng/mL human recombinant FN1, (**k**) T98-G, (**l**) LN-229, (**m**) U87-MG, and (**n**) U-118 MG. (**o-p**) Transwell invasion results for glioblastoma cell lines exposed to human recombinant FN1, (**o**) T98-G and (**p**) LN-229. Data were generated from n=3 independent experiments and displayed as mean ± S.D. One-way ANOVA: * < 0.05, ** < 0.01, *** < 0.001, and **** < 0.0001.

To directly assess this, we exposed the T98-G, LN-229, U87-MG, and U-118 MG cells to human recombinant FN1s (rFN1) of different concentrations. Indeed, rFN1 induced the migration of all four cell lines (**Figure 4k-n**). For invasion, it induced the invasion of T98-G and LN-229 (**Figure 4o, p**) but not U87-MG and U-118 MG cells (**Figure S5a-c**). Thus, we conclude that FN1 is a key ligand expressed by CAFs that can induce malignant cell migration and invasion in glioblastoma *in vitro*.

### CAFs are associated with a proneural-to-mesenchymal transition in glioblastoma

Proneural-to-mesenchymal transition (PMT), a molecular process analogous to epithelial-to-mesenchymal transition (EMT), contributes to malignant cell plasticity in glioblastoma (*33*). In addition, PMT is often associated with enhanced cellular motility and invasion (*33*), two phenotypes observed above when malignant cells were exposed to CAFs *in vitro*. Thus, we sought to evaluate if CAFs are involved in PMT. Analysis of the TCGA and CGGA cohorts indicated that glioblastoma tumors with a high CAF enrichment score were significantly associated with higher activities of the EMT hallmark gene set (**Figure 5a**). Furthermore, TCGA glioblastomas with a high CAF score were enriched for several independent gene sets describing the GBM mesenchymal subtype (i.e., Verhaak_Mesenchymal, Phillips_Mesenchymal, Neftel_MES_Module, and Wang_mGSC) (*39-42*) (**Figure 5b**). In contrast, TCGA glioblastomas with a low CAF score were enriched for gene sets associated with the proneural subtype (i.e., Verhaak_Proneural, Neftel_NPC_module, Neftel_OPC_Module, and Phillips_Proneural) (**Figure 5b**). Interestingly similar results were observed upon analysis of both the CGGA 325 and CGGA 693 datasets (**Figure S6a, b**). To study this in more detail, PCA was performed on TCGA glioblastomas classified as either proneural or mesenchymal subtype, first stratified by subtypes and then by CAF scores to generate four groups (i.e., proneural CAF low/high and mesenchymal CAF low/high). This analysis showed that these glioblastoma tumors resided on a single axis of continuous variation in which the proneural CAF low and the mesenchymal CAF high were enriched on the two opposite ends of the PC1 axis, with the other two groups between (**Figure 5c**). Similar results were observed from the analysis of the CGGA 325 cohort (**Figure S6c**). Interestingly, as one moved along the PC1 axis from proneural CAF low to mesenchymal CAF high, there was a gradual increase in the expression of genes related to inflammation, invasion, and the mesenchymal subtype and a corresponding decrease in the expression of genes related to the proneural subtype, in both the TCGA (**Figure 5d**) and the CGGA 325 dataset (**Figure S6d**). Collectively, these results indicate that CAFs are strongly associated with the mesenchymal-subtype and moreover the activation of the CAF-related gene programs is highly correlated with the transition from a proneural- to a mesenchymal-state.

**Figure 5.**
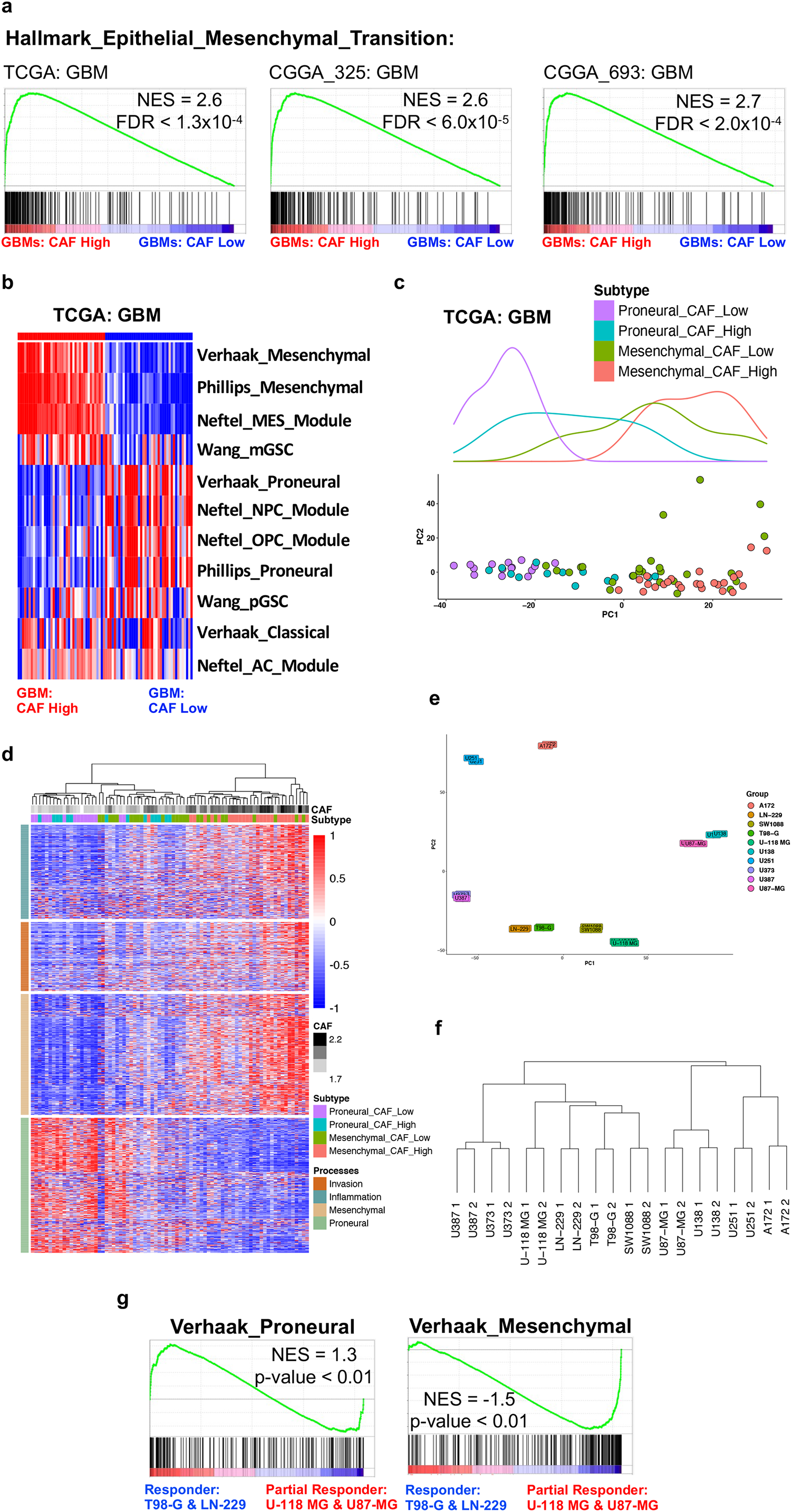
CAFs are associated with a proneural-to-mesenchymal transition in glioblastoma. (**a**) GSEA enrichment plots for the Hallmark epithelial mesenchymal transition gene set, comparing glioblastomas with a high to low CAF enrichment score in the TCGA and CGGA datasets. (**b**) Heatmap depicting individual enrichment scores for signature gene sets describing specific glioblastoma subtypes among glioblastoma samples (columns) with a high and low CAF enrichment score in the TCGA dataset. (**c**) PCA plot for glioblastomas samples in the TCGA cohort, with colors for four groups: proneural CAF low (n=15), proneural CAF high (n=14), mesenchymal CAF low (n=25), and mesenchymal CAF high (n=25). The densities of samples are shown above. (**d**) Heatmap showing changes in the expression of genes associated with inflammation, invasion, mesenchymal-subtype, and proneural-subtype as glioblastomas accumulate more CAFs in the TCGA cohort. (**e**) PCA plot of microarray data for glioma cell-lines (n=2 samples for each cell line). (**f**) Dendrogram showing correlation of microarray samples from glioma cell-lines. (**g**) GSEA enrichment plots for the proneural and mesenchymal glioblastoma subtype gene sets, comparing responder cell lines (T98-G and LN-229) and partial responder cell lines (U118-MG and U87-MG).

As stated above, the effects of Fibro^CM^, CAF^CM^, and rFN1 on cell migration and invasion varied among the four cell lines. Fibro^CM^, CAF^CM^, and rFN1 consistently induced the migration and invasion of the T98-G and LN-229 cells, but they conferred variable effects on the migration and invasion of the U87-MG and U-118 MG cells (**Figure S5c**). As such, T98-G and LN-229 cell lines were labeled as *responders* while U87-MG and U-118 MG cell lines as *partial responders*. PCA and clustering analysis of the microarray data from 10 glioma cell lines (*43*) indicated that the *responders* were clustered together but separately from the *partial responders* (**Figure 5e, f**). The distinction was also observed using RNA-seq data from the cancer cell line encyclopedia (CCLE) database (**Figure S6e**). Lastly, GSEA of the microarray data suggests that the *responders* and *partial responders* were enriched for gene sets describing the proneural- and mesenchymal-subtype, respectively (**Figure 5g**). Similar enrichment trends were observed when the CCLE RNA-seq data were analyzed (**Figure S6f)**.

Collectively, these results suggest that glioblastoma cells in a proneural-like state could be more likely to migrate and invade upon exposure to CAF^CM^, compared to glioblastoma cells in a mesenchymal-like state. In addition, these results raise the possibility that malignant cells in the proneural-like state are likely to adopt mesenchymal-like characteristics (i.e., enhanced migration and invasion) upon exposure to CAF^CM^; thus, indicating that CAFs are potentially associated with promoting PMT (see Discussion).

### CAFs are enriched in perinecroti/pseudopalisading zones and in the same vicinity of malignant mesenchymal cells

The observation that CAFs are highly associated with the mesenchymal subtype and might even facilitate PMT prompted us to study if CAFs are located closely to mesenchymal malignant cells in the actual GBM tumors. For this, we examined RNA-seq data describing anatomical regions in human glioblastoma samples that was acquired by the IVY Glioblastoma Atlas Project (IVY GAP) (*25*). We compared the CAF enrichment scores among different regions and found that CAFs were highly enriched in perinecrotic and pseudopalisading zones but relatively depleted in the leading edge of the tumor (**Figure 6a**). Further examination of the *in-situ* hybridization data (*25*) demonstrated that several CAF maker genes from our scRNA-seq analysis were detected in the perinecrotic/pseudopalisading zones across several patients, such as *ITGA5, COL1A1, CAV1*, and *ANGTPL4* (**Figure 6b**). Furthermore, in multiple patient samples, *FN1* was also expressed in the perinecrotic and pseudopalisading zones (**Figure 6c**). Additionally, genes expressed higher in the perinecrotic/pseudopalisading zones were enriched for gene sets describing the mesenchymal subtype but not the proneural subtype (**Figure 6d**). Collectively, these results indicate that CAFs are primarily enriched in perinecrotic/pseudopalisading zones, with the neighbor malignant cells likely to be in the mesenchymal state.

**Figure 6.**
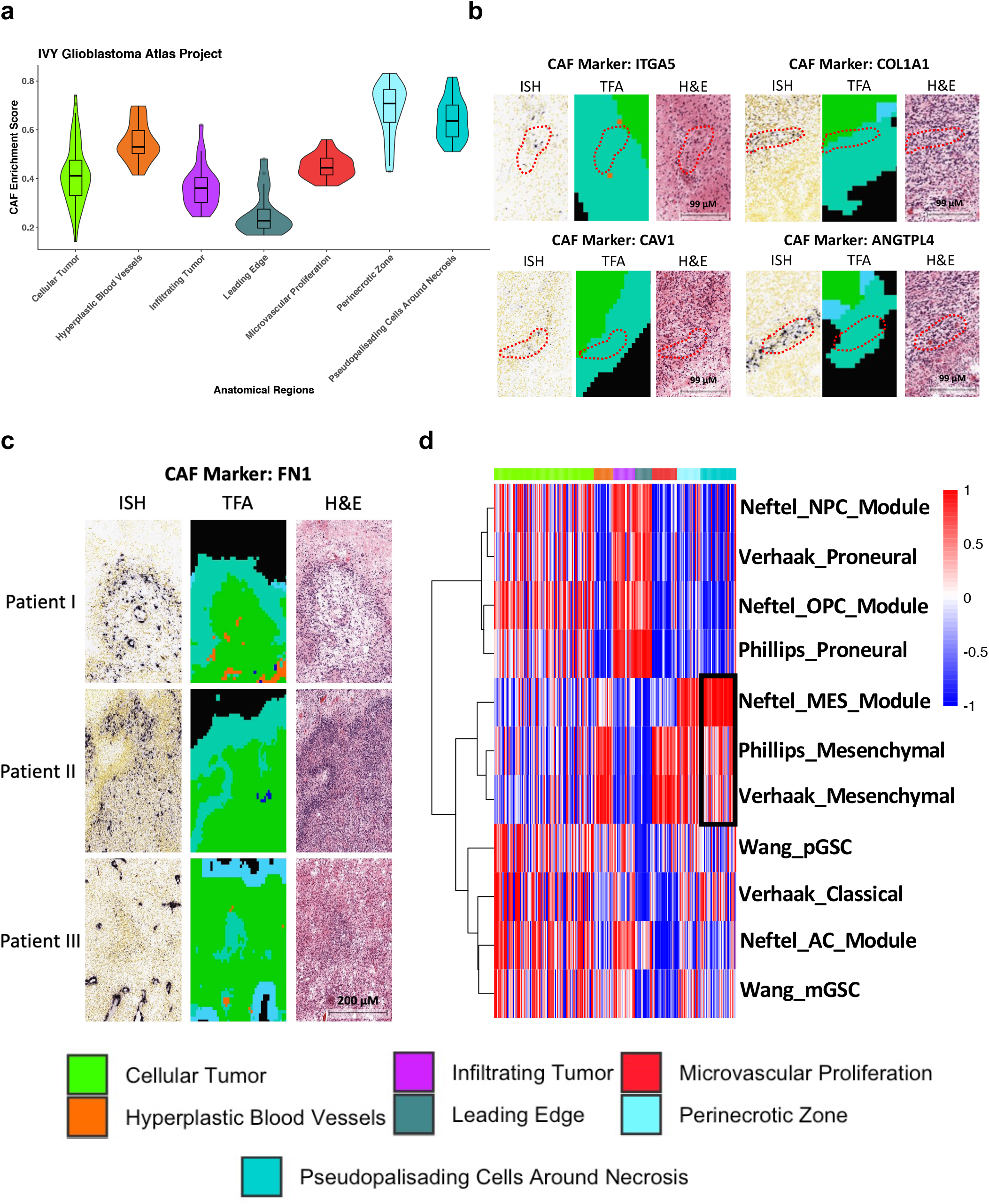
CAFs are enriched in the perinecrotic/pseudopalisading zones and in the same vicinity of malignant mesenchymal cells. (**a**) Violin plot of CAF enrichment scores across anatomical regions in the IVY-GAP atlas. (**b**) *In situ* hybridization expression in several human glioblastoma tissue samples for selected CAF marker genes (*ITGA5, COL1A1, CAV1*, and *ANGTPL4*) in the perinecrotic and pseudopalisading zones. For each marker there are three plots: *in situ* hybridization staining (ISH), tumor feature annotation (TFA) indicating which anatomical region the marker gene is expressed, and hematoxylin and eosin staining (H&E). Scale bars, 99 µm. (**c**) *In situ* hybridization expression analysis for *FN1* in several human glioblastoma samples. Scale bars, 200 µm. (**d**) Heatmap of enrichment scores for signature gene sets describing glioblastoma subtypes across anatomical regions in the IVY-GAP atlas.

## Discussion

In this study, we have focused on identifying CAFs, characterizing their signature genes, and studying their interactions with glioblastoma malignant cells. We did not find genes uniquely expressed in CAFs, indicating a difficulty in applying one or two genes to specifically label CAFs in GBM tumors. Our current findings build upon several studies suggesting that CAFs could be an important cell-type mediating glioblastoma pathogenesis. Specifically, Clavreul *et al* isolated a population of cells termed glioblastoma-associated stromal cells (GASCs), which highly expressed several CAF markers (i.e., ACTA2 and PDGFRB) and contractile markers (i.e., CNN1, MYH11, and PFN1), suggesting GASCs have a hybrid CAF/myofibroblast phenotype (*44*). Intracerebral co-injection of U87-MG cells with GASCs showed an increase in small vessel development compared to control models, suggesting that GASCs could be involved in mediating angiogenesis (*44*). Others have shown that conditioned media harvested from CAF cells established from skin metastases of primary nodular melanoma enhanced the chemotaxis of U87-MG cells (*45*). However, it is not clear how CAFs from metastatic skin tumors are comparable to GBM-derived CAFs. Furthermore, that study did not identify the CAF-secreted factors mediating the enhanced migratory phenotype (*45*). Recently, scRNA-seq analysis of IDH mutant and wild type (WT) gliomas reported the presence of fibroblasts, based on the expression of *DCN* and *FBLN1* (*46*). Consistently, our scRNA-seq analysis identified *DCN* as a marker gene for CAFs (but not specific) and DCN was also detected in the CAF-secretome. Collectively, these studies support that CAFs are present in glioblastoma; however, how they mediate pathogenesis remained not established.

By contrast, our study provides a broader investigation of the molecular features and functional roles of CAFs in glioblastoma. Consistent with the literature on CAFs in epithelial tumors (*22*), we showed that glioblastoma derived CAFs facilitate the migration and invasion of malignant cells. Mechanistically, our results indicate that CAF-derived FN1 contributes to the enhanced migratory and invasive phenotype. Other studies have also found the importance of FN1 in glioblastoma. For example, GBP2-promotion of glioblastoma cell invasion is dependent on STAT3 signaling and FN1 induction (*36*). In addition, others have suggested that FN1 expression promotes glioblastoma cell invasion of the basement membrane and loss of FN1 leads to reduced tumor growth and angiogenesis (*37*). Our study indicates that CAFs are a key source for FN1 in glioblastoma and thus targeting CAFs could potentially reduce FN1-induced tumorigenesis, specifically malignant cell invasion.

Analysis of bulk human tumors indicates that glioblastomas can be categorized into three groups with unique gene programs. Pioneering work by Phillips *et al* described the proliferative, proneural, and mesenchymal subtypes (Phillips subtypes) (*40*). While other studies defined glioblastoma into three transcriptomic subtypes: proneural, mesenchymal, and classical (Verhaak subtypes) (*41, 47*). Among them, the derivation of the mesenchymal subtype is the least understood. It is known that some genetic alterations, such as NF1 mutation or deletion, are enriched among mesenchymal GBMs. Given that mesenchymal state/subtype is associated with poor treatment response and a slightly worse clinical outcome, delineating the underlying mechanisms driving the establishment of mesenchymal glioblastoma remains an interesting area. Cell-to-cell interaction within TME is certainly a key proponent to driving a mesenchymal-like state in glioblastoma. Multiple studies have demonstrated that macrophages are highly associated with the mesenchymal subtype (*3, 4*). Recently, Hara *et al* showed that macrophage-derived oncostatin M (OSM) drives a mesenchymal-like state by inducing STAT3 signaling through interreacting with either OSMR or LIFR in complex with GP130 on glioblastoma cells (*4*). Data from this study suggest that glioblastomas with a high CAF enrichment score were significantly correlated with several independent gene sets describing the mesenchymal state in glioblastoma, indicating that, like macrophages, CAFs are associated with the mesenchymal subtype. In this case, the association seems to be mediated by CAF-derived FN1, as FN1 can enhance two hallmarks associated with the mesenchymal phenotype: cell motility and invasion. In the future, it will be interesting to study if soluble FN1 can increase the expression of a mesenchymal program, thus directly supporting the hypothesis that TME interactions involving CAFs can assist in establishing mesenchymal glioblastomas.

Recent scRNA-seq analysis of glioblastoma patient samples indicated that malignant cells themselves (vs bulk tumors discussed above) could reside in four distinct cellular states: neural progenitor-like (NPC-like), oligodendrocyte progenitor-like (OPC-like), astrocyte-like (AC-like), and mesenchymal-like (MES-like) (Neftel subtypes) (*39*). Its NPC- and OPC-like states are correlated with the Verhaak proneural subtype, while the MES-like state with the Verhaak mesenchymal subtype (*39*). Functional study further suggests that malignant cells can transition between states and that the state diversity is maintained through cellular plasticity (*39*). These further underscores transcriptomic plasticity as a key issue in glioblastoma that mediates therapeutic resistance, sustained growth, and enhanced invasiveness (*3, 39*). The underlying process were referred in literature as proneural-to-mesenchymal transition, analogous to EMT, in which environmental cues causes a transition from a proneural- to a mesenchymal-transcriptomic state in glioblastoma cells (*33*). Other studies have shown that CAFs can mediate EMT in other cancer types (*48*). Interestingly, in this study, we not only observed that glioblastomas with a low CAF enrichment score were highly associated with gene sets describing the proneural state, but also showed that as glioblastomas acquired CAF gene program there was a gradual decrease in the expression of genes related to the proneural subtype while a corresponding increase in the expression of genes associated with inflammation, invasion, and the mesenchymal subtype. In addition, this study suggests that *in vitro* glioblastoma cells resembling the proneural-state seem to respond better to the CAF cues than cell lines resembling the mesenchymal-state, raising the possibility that CAFs may have distinct functions at different states of the PMT. Taken together, these results suggest that CAFs could be another microenvironmental factor contributing to glioblastoma cell plasticity, specifically by facilitating PMT. Future studies, however, would be needed to evaluate if CAF-derived FN1 or other factors in the CAF-secretome can directly mediate PMT.

We focused on FN1 in the current report, but it is important to note that several other proteins identified in our CAF-secretome analysis could also be of importance. For example, several collagen factors such as COL1A1, COL1A2, and COL6A1 were abundantly detected. Depletion of myofibroblast derived collagen I was associated with immunosuppression in pancreatic cancer (*49*), suggesting that CAF-derived collagen could act as a potential barrier mediating immune-cell trafficking. SERPINE1, detected in the CAF-secretome, is also very interesting, because studies have indicated that it is involved in glioblastoma cell invasion (*50*) and moreover CAF-derived SERPINE1 has been directly implicated in esophageal squamous carcinoma cell migration and invasion (*51*). In addition, although our study focused on the crosstalk between CAFs and malignant cells, in reality TME is a dynamic system involving the interactions of many different stromal cell types. Thus, in the future it would be interesting to explore how CAFs interact with other cell types and collectively affect glioblastoma pathogenesis.

How CAFs are derived and maintained in glioblastoma is an open question, especially since it is thought that healthy brain tissue lacks an abundance of fibroblasts. However, *in vitro* models suggest that conditioned media harvested from U87-MG cells induced the expression of several myofibroblast related markers in human mesenchymal stem cells (hMSCs), alluding to hMSCs as a potential cellular source for CAFs in glioblastoma (*52*), which is partially supported by recent cell lineage tracing analyses (*52*). Understanding what cell types or environmental factors contribute to CAF accumulation in glioblastoma could lead to novel therapeutic strategies that either suppress CAF activation or directly deplete the CAF population in the TME. Results presented in this study suggest that CAFs are highly enriched in the perinecrotic and pseudopalisading zones in the glioblastoma. It would be informative to assess if hypoxia is sufficient to induce CAF activation and stabilization. In addition, it was observed that mesenchymal-like cells are also highly enriched in the vicinity. CAF interactions with surrounding malignant cells thus might assist in glioblastoma cell adaptation to hypoxic regions. It would be interesting to evaluate if FN1 plays a role there, since *in situ* hybridization data suggests that it is also expressed in the same zones. Understanding these various mechanisms would be important for broadening the current understanding of how CAFs mediate pathogenesis in glioblastoma.

In summary, this study proposes the following model. In proneural glioblastomas there is a low abundance of CAFs, however, as glioblastomas gradual accumulate CAFs within the microenvironment they become more mesenchymal (**Figure 7**). The accumulation of CAFs contributes to soluble FN1 and other cues, which in turn will contribute to an enhanced migratory and invasive phenotype among malignant cells (**Figure 7**). This enhancement ultimately contributes to a higher tumor grade and a shorter overall survival (**Figure 7**). Overall, this study highlights the importance of studying CAFs and their molecular functions in mediation of glioblastoma pathogenesis, thus, alluding to a novel therapeutic target in glioblastoma.

**Figure 7.**
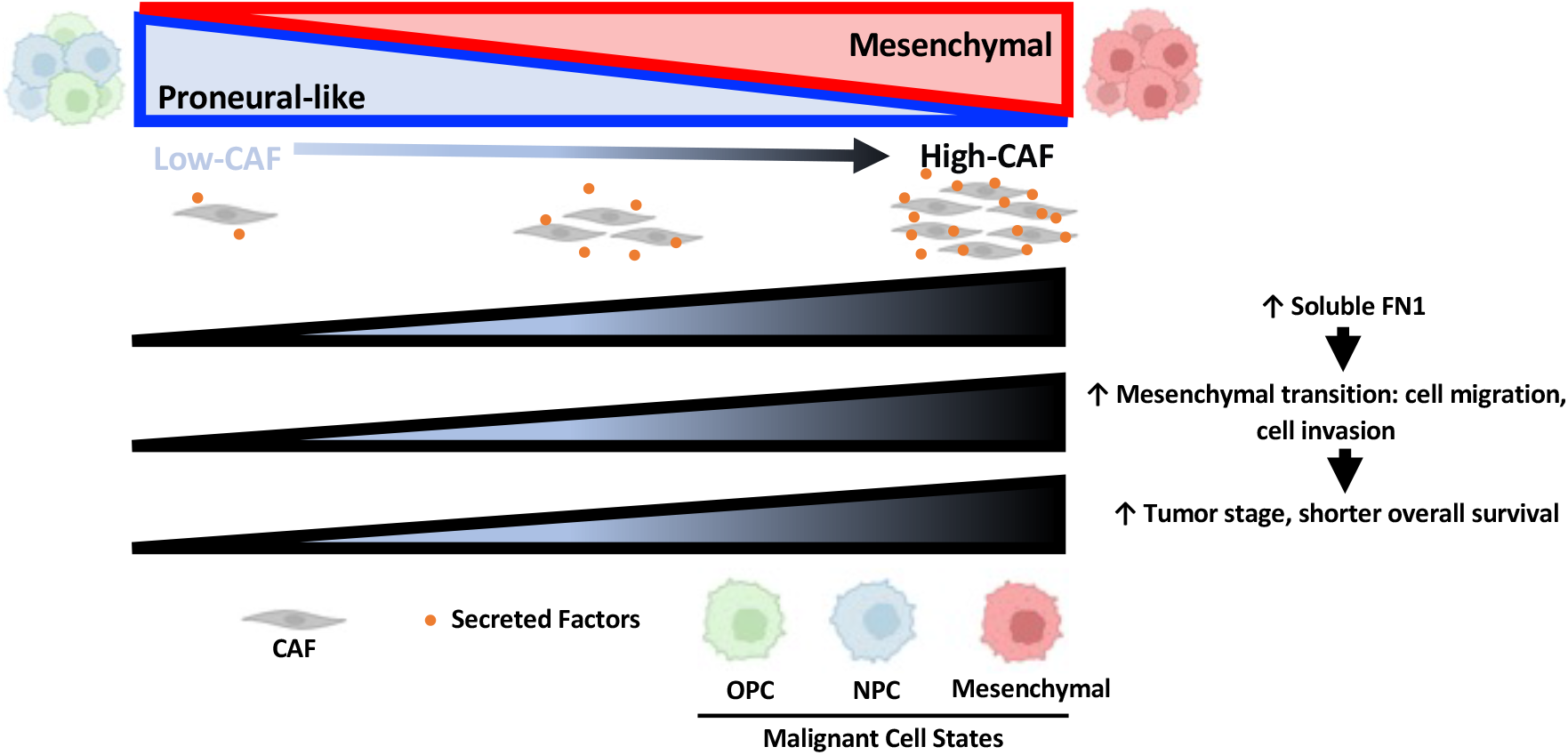
Working model of CAF contribution in glioblastoma.

## METHODS

### Datasets

The Cancer Genome Atlas (TCGA) low grade glioma (LGG) and glioblastoma bulk RNA-seq datasets with normalized read counts and FPKMs (fragment pet kilobase of transcript per million mapped reads) were downloaded from the UCSC Xena platform (*19, 53*). Similar data were downloaded for CGGA 325 and CGGA 693, from the Chinese Glioma Genome Atlas (CGGA) (*18*). Deidentified patient data corresponding to the samples were also downloaded. Normalized scRNA-seq data for GBM samples (described below) was acquired from the UCSC cell browser (cells.ucsc.edu), while the raw count data was generously provided by the lab of Arnold Richard Kriestein (*17*). Count and RPKM (reads per kilobase of transcript, per million mapped reads) RNA-seq data for glioblastoma cell lines were downloaded from the cancer cell line encyclopedia atlas (CCLE) (*54*). Microarray data for glioma cell lines was downloaded from the gene expression omnibus (GEO; accession number GSE4536) (*43*). RNA-seq data from anatomic structures of glioblastoma was downloaded from the IVY glioblastoma atlas project (IVY-GAP) (*25*).

### Estimating the Proportions of Immune and Cancer Cells (EPIC) in glioblastoma tumors

Cell type proportions in the bulk RNA-seq samples of low grade glioma (LGG) and glioblastoma from the TCGA (*19*) and CGGA (*18*) were calculated using EPIC (v1.1.5) (*20*). Using the *EPIC* function and with *refences* set to “TRef”, proportions of B Cell, CD4+ T Cell, CD8+ T Cell, NK Cell, macrophage, endothelial, and CAF were estimated for each sample in the TCGA (*19*) and CGGA (*18*) datasets. Next, using STATA/IC 15.1, survival analysis was first assessed with a univariate cox proportional hazards model, with proportions of individual cell types coded as a continuous variable. Kaplan-Meier estimator method was used to compare patients that were divided into quartiles based on the calculated CAF proportions. Quadrant 1 (tumors with the highest estimated CAF proportion) was compared with quadrant 4 (tumors with the lowest estimated CAF proportion). A log-rank and HR test were used to determine statistical significance between the two groups of patients, as previously described (*55*).

### Glioblastoma scRNA-seq analysis

Single-cell RNA sequencing data describing 32,877 cells dissociated from 11 primary glioblastoma resections was obtained from Bhaduri *et al (17*). The authors characterized cell types as oligodendrocyte precursor cells (OPCs), microglia, tumor-associated macrophages, several populations of neurons, radial glia, glial cell populations with varying degrees of maturity, and malignant cells (*17*). This information was provided as metadata and directly used in our analysis. CAFs in these data, however, were identified by us using the *findoutliers* function in RaceID (v0.1.9) on 10,000 randomly selected non-malignant cells (*21*). Specifically, the probability threshold was set to 1×10^−3^, minimal transcript count was set to 5, minimal number of outlier genes required was set to 2, and cells were merged to outlier clusters if their distances were smaller than 0.95 (default). Differential expression analysis was performed by comparing cells in the CAF cluster to all other non-malignant cells-types using the *clustdiffgenes* function in RaceID (*21*). A gene was considered significantly upregulated in the CAF cluster if its expression difference reached a log2-(fold change) > 2 at adjusted p value < 0.05. Such genes were considered as CAF markers or signature genes. They were screened against 1,368 scRNA-seq datasets in the PanglaoDB (*23*) to identify cell types that predominantly express these genes. Cell-to-cell interactions across the cell types was assessed using CellChat (v1.1.3) (*56*). Specifically, CellChat was used to predict ligand-receptor interactions by analyzing the mean expression for genes that correspond to ligands and receptors in a curated ligand-receptor database. Receptors that correspond to the FN1 ligand was specifically homed in on to determine how FN1 from CAFs could interact with malignant cells and other TME cells.

### Spatial Transcriptomics Analysis

Human glioblastoma whole transcriptome, visium spatial dataset by the Space Ranger (1.2.0; 10x Genomics) was downloaded from the 10x Genomics website on March 5, 2021. Analysis and visualization of the spatial dataset were performed using Seurat (v4.0.4) (*57*) with distinct transcriptomic regions determined by the *FindClusters* function, using resolution 0.6. Expression of individual genes was visualized using the *SpatialFeaturePlot*. The *FindMarkers* function was used to perform differential gene expression analysis with a Wilcoxon rank-sum test to define marker genes for each region, using the default parameters. To identify CAF enriched regions, MuSiC (v0.2.0) (*24*) was utilized to deconvolute detected spots in the spatial dataset and to estimate cell-type-specific abundances via the *music_prop* function (default parameters). Estimate CAF-type abundances were projected as a density heatmap on the glioblastoma spatial plot using ggplot2 (v3.3.5).

### Gene set variation analysis (GSVA)

The top 100 marker genes for our newly detected CAF population and each of the other cell types described by Bhaduri A *et al* for the same glioblastoma scRNA-seq data (*17*) were utilized by GSVA (v1.40.1) (*58*) to calculate cell-type enrichment scores for each of the LGG and glioblastoma samples in the TCGA (*19*), CGGA (*18*), and IVY-GAP (*25*) datasets. Specifically, *gsva* function was used with RNA-seq count data and the kcdf parameter set to ‘possion’. The association between cell-type enrichment scores and clinical outcomes was first assessed with a univariate cox proportional hazards model with the enrichment scores coded as a continuous variable in STATA/IC 15.1. As the analysis described above, this was followed up with Kaplan-Meier estimator in which patients were divided into quartiles based on the calculated CAF enrichment scores. Quadrant 1 (glioblastomas with the highest CAF enrichment scores) was compared with quadrant 4 (glioblastomas with the lowest CAF enrichment scores). A log-rank and HR test were used to determine statistical significance between the two groups. The association of CAF enrichment scores to clinical outcome in LGG and glioblastoma patients was also assessed with a multivariate cox proportional hazards model using STATA/IC 15.1. For LGG datasets, the impact of the CAF enrichment scores was assessed while adjusting for variables including age (continuous), gender (dichotomized), IDH1 mutation status (dichotomized), and neoplasm grade (continuous). For glioblastoma datasets, the variables included age (continuous), gender (dichotomized), IDH1 mutation status (dichotomized), and O-G-methyl guanine-DNA methyl transferase (MGMT) methylation status (dichotomized). In the TCGA glioblastoma cohort, the association between CAF enrichments scores and disease-free progression was assessed with a Kaplan-Meier estimator analysis, in which patients were stratified into two groups based on the median CAF enrichment score. A log-rank and HR test were used to determine statistical significance between the two groups of patients. This analysis was not performed on the CGGA datasets because their clinical information did not document disease-free progression.

### Comparing glioblastomas with high and low CAF enrichment scores

Principal component analysis and differential gene expression analysis comparing glioblastomas with a high and low CAF enrichment score (derived from GSVA) in the TCGA and CGGA cohorts were performed using DESeq2 (v1.32.0) (*59*). Gene set enrichment analysis (GSEA, v4.1.0) (*30*) was performed to identify gene sets enriched in genes expressed higher in glioblastomas with high (or low) CAF enrichment scores. Specially, genes were ranked by multiplying the -log_2_ transformation of the p value with the sign of the log_2_ (fold change). Hallmark, REACTOME gene sets, and Oncogenic Signatures gene sets in the Molecular Signatures were analyzed, with the enriched sets (q-value < 0.05) selected for further analysis using the enrichment map plug-in for cytoscape (v3.8.2) (*31, 60*).

### Cell lines and cell culture

Human glioblastoma cell lines (i.e., LN-229, U87-MG, T98-G, and U-118 MG) were cultured under standard condition at 3°C in DMEM supplemented with 10% FBS and 1% penicillin and streptomycin under a humidified atmosphere of 5% CO_2_/95% air. T98-G and IMR-90 cell lines were cultured at 3°C EMEM supplemented with 10% FBS and 1% penicillin and streptomycin under a humidified atmosphere of 5% CO_2_/95% air. Glioblastoma-derived CAFs were purchased from the Vitro Biopharma (catalog number: CAF03). They were cultured at 3°C in MSC-GRO VitroPlus III Low Serum Complete Medium (Vitro Biopharma catalog number: PC00B1) under a humidified atmosphere of 5% CO_2_/95% air.

### Immunocytochemistry

Glioblastoma-derived CAFs were fixed by adding 1mL with 10% formalin for 20 minutes at room temperature. The plate was washed three times with 1x PBS. 1mL of 0.1% triton-100 was add for 10 minutes at room temperature to permeabilize the cells. The cells were then washed again three times with 1x PBS. 5% goat serum was added and incubated with the cells for 1 hour at room temperature. Then either α-SMA (Sigma Aldrich, Catalog Number: A5228), Vimentin (Santa Cruz Biotechnology Inc, Catalog Number: sc-373717), or GFAP (Santa Cruz Biotechnology Inc, Catalog Number: sc-166458) at a 1:100 dilution was added and incubated overnight at 2-8°C. The next day, cells were washed three times with 1x PBS. Secondary antibody (Alexa Fluor 488-conjugated AffiniPure Goat Anti-Mouse IgG) at concretion of 1:1000 diluted in 1mL of 1x PBS was added to cells. Cells were incubated with secondary antibody for 1 hour at room temperature on a shaker. Cells were washed again three times with 1x PBS. Nuclei of cells were stained with 25ug/mL Hoecst 33258 for 10 minutes at room temperature. The cells were washed a final three times with 3 mL of 1x PBS and read on a fluorescent microscope. Images from immunocytochemistry analysis was provided by Vitro Biopharma.

### Preparation of conditioned medium

5×10^5^ IMR-90, or glioblastoma CAFs cells were seeded in 100 mm dishes in cell culture conditions described above. After 24 hours, the media was aspirated, and the cells were gently washed with 5 mL of PBS. Then cells were cultured for additional 72 hours in serum-free DMEM. The media was collected, centrifuged (500g, 20°C, 5 minutes), and immediately used for cell migration and invasion assays.

### Transwell migration and invasion assay

4×10^4^ of either the T98-G, LN-229, U87-MG, and U-118 MG cell lines were seeded in the top chamber of a 24 well transparent PET membrane with 8.0 uM pore size (Falcon reference number: 353097). 750ul of either fibroblast conditioned media (Fibro^CM^) or CAF conditioned media (CAF^CM^) were added to the bottom chamber. Cell lines were allowed to migrate for 24 hours at standard cell culture conditions described above. Following the migration incubation period, cells were then fixed and stained with a solution containing methanol 5% (vol/vol), crystal violet 0.5% (wt/vol) in H_2_O. Five images were captured per transwell at 10x magnification and the number of cells per field were then counted using ImageJ (*61*). When testing the effects of human recombinant FN1 (rFN1) (R&D Systems Catalog number: 4305-FNB), rFN1 was prepared in serum free DMEM at the concentration of either 10 ng/mL or 20 ng/mL. As a negative control, all conditions were compared to cells exposed to serum free DMEM. For invasion assays a Matrigel invasion chamber (corning reference number: 354480) was used.

### Protein extraction and digestion

A speedVac centrifuge was used to dry 1 ml of cell media. Disulfide bond reduction was then performed by resuspending the dried samples in a buffer containing 5% SDS, 5 mM DTT, and 50 mM ammonium bicarbonate (pH = 8) and left on the bench for roughly 1 hour. Samples were then alkylated with 20 mM iodoacetamide in the dark for 30 minutes. Afterward, phosphoric acid at a final concentration of 1.2% was added to each sample, followed by dilution in six volumes of binding buffer (90% methanol and 10 mM ammonium bicarbonate, pH 8.0). After gentle mixing, the protein solution was centrifuged at 500 g for 30 sect following loading into a S-trap filter (Protifi). Next, 1 µg of sequencing grade trypsin (Promega) diluted in 50 mM ammonium bicarbonate was added to the S-trap filter. Samples were then digested for 18 hours a 3°C. Peptides were then eluted by first adding 40 µl of 50 mM ammonium bicarbonate followed by 40 µl of 0.1% TFA and concluded with the addition of 40 µl of 60% acetonitrile and 0.1% TFA. The peptide solution was then pooled together and centrifuged for 30 seconds at 1,000 g. Lastly, the peptide solution was then dried in a vacuum centrifuge.

### Nano liquid chromatography coupled to online mass spectrometry

Samples were loaded onto a Dionex RSLC Ultimate 300 (Thermo Scientific) coupled online with an Orbitrap Fusion Lumos (Thermo Scientific) following resuspension with 10 µl of 0.1% TFA. A C-18 trap cartridge (300 µm ID, 5 mm length) and a picofrit analytical column (75 µm ID, 25 cm length) packed in-house with reversed-phase Repro-Sil Pur C18-AQ 3 µm resin was used to perform chromatographic separation. At a flow rate of 300 nl per minutes the peptides were separated using a 120-minute gradient from 4-30% buffer B (buffer A: 0.1% formic acid, buffer B: 80% acetonitrile + 0.1% formic acid). Mass spectrometer was then set to acquire spectra in a data-dependent acquisition (DDA) mode. The full mass spectrometer was set to 300-1,200 m/z in the orbitrap with a resolution of 120,000 (at 200 m/z) and an AG target of 5×10e5. Last via top speed mode (2 seconds), HCD collision energy of 35, and a AGC target of 1×10e4 setting were used in the ion trap when performing MS/MS.

### Proteome data processing, statistical analysis, and data deposition

Proteome Discoverer software (v2.4, Thermo Scientific) using SEQUEST search engine and the SwissProt human database (updated February 2020) was used to search the raw proteome data. The total proteome search encompassed variable modification of both fixed modification of carbamidomethyl cysteine and N-terminal acetylation. With the digestive enzyme specified as trypsin and 2 missed cleavages were allowed. 10 pm for precursor ions and 0.2 Da for-product ions were the parameters for mass tolerance with peptide and protein false discovery rate were set to 1%. Data was log2-transformed and the average distribution was used to normalize data with missing value imputed as previously described (*62*). Raw data are deposited on the publicly available repository Chorus (https://chorusproject.org) under the project number #1754.

### Protein rankin

To identify a list of proteins that could mediate the observed migration and invasion phenotype, a ranking method was developed. The variables used in ranking included: if a protein was detected in both the Fibro^CM^ and CAF^CM^, protein abundance in the samples, expression difference of the encoding gene in the CAF cluster vs other cell types in the glioblastoma scRNA-seq data, Pearson correlation coefficient for the encoding gene’s correlation with CAF enrichment score calculated by GSVA using the TCGA, CGGA 325, and CGGA 693 datasets, and hazard ratios for the encoding gene in the survival analysis. In addition, a positive weight was added if the encoding gene was expressed in the CAF enriched region in the spatial transcriptomic dataset or in the IVY-GAP atlas, if the hazard ratios were significant (p-value < 0.05), and if the Pearson correlation coefficients was significant (p-value < 0.05). The scores from each of the variables were summed for ranking proteins. Lastly, among the top ranked proteins, the availability of human recombinant protein was considered so that functional validation experiments could be conducted.

### Proteomic data integration

The limma package (v3.48.3) (*63*) was used to perform integrated analysis of the whole cell lysate (WCL) proteomic data generated from glioblastoma-derived CAF cell lines, CAFs isolated from human oral tongue squamous cell carcinoma (OTSCC) (ftp://massive.ucsd.edu/MSV000081585) (*34*), and resting and activated CD4+ T Cells isolated from two separate donors (http://proteomecentral.proteomexchange.org) (*35*). Specifically, batch effects across datasets were removed using the *removeBatchEffect* function in limma. After batch effects were removed, PCA was done using the *prcomp* function and Pearson correlation across samples was calculated using the *cor* function and then used to cluster samples with corrplot (v0.90).

### Statistics and reproducibility

The sample size was based on the number of samples available for all human samples. For whole cell lysate and secretome proteomic analysis of both the glioblastoma-derived CAF and IMR-90 cell lines, two replicates were used. There was no data exclusion from any experiment in our analysis. Heatmaps and unsupervised hierarchical cluster analysis, using Euclidean distance measurements, were performed using the pheatmaps package in R. Dendrogram analysis using Euclidean distance measurements was performed using the corrplot package and clustered using ward.D2 method in the hclust function in R. Pearson Correlation analysis was performed using the cor.test function in R. *In vitro* experiments, where indicated, were conducted in either three or four independent experiments and displayed as mean ± S.D. Presentation and statistical analyses of *in vitro* cell experiments were performed using GraphPad Prism software. Statistical significance for assays were determined using either a t test of one-way analysis of variance (ANOVA) assuming normal distribution and equal variance. An observation was considered statically significant at p < 0.05.

## Data and code availability

All the scRNA-seq, bulk RNA-seq, and spatial transcriptomics data were from either previously publications or public domains. The code for the analysis will be made available at the GitHub (https://github.com/bioinfoDZ).

## Acknowledgements

We would like to thank Vitro Biopharma for the immunocytochemistry analysis of the GBM-CAF and the Kriestein lab for sharing the GBM scRNA-seq data. This work is supported by the NIH/NCI (R01CA255643, R01CA262903, and R01CA259480, to D.Z. and R01CA175495 to X.Z.), and U.S./DOD (BC190403, to X.Z.). P.M.G. is supported by the NIH/National Center for Advancing Translational Science (NCATS) Einstein-Montefiore CTSA training grant (TL1 TR002557). S.S. would like to acknowledge the supports from AFAR (Sagol Network GerOmics award), Deerfield (Xseed award), Relay Therapeutics, Merck, and the NIH Office for the Director (1S10OD030286-01).

## Authors’ Contributions

**P.M.G**.: Conceptualization, experimental design and execution, data curation, method development, formal analysis, statistics, writing original draft, and editing the manuscript. **Y.L**.: data curation and method development. **M.P**.: conceptualization. **Y.W**. and **A.T.M**.: experimental design. **S.G**. and **S.S**.: generation and analysis of proteomic data. **C.M**. and **J.S**.: experimental design. **X.Z**.: Experimental design, supervision, funding acquisition, and manuscript editing. **D.Z**.: Conceptualization, experimental design, supervision, funding acquisition, writing manuscript, project administration.

## Declaration of Interest

The authors declare no competing interests.

## Notes

### Competing Interest Statement

The authors have declared no competing interest.

