## supplemental figures & Tables for "Functional Contribution of Cancer-Associated Fibroblasts in Glioblastoma"

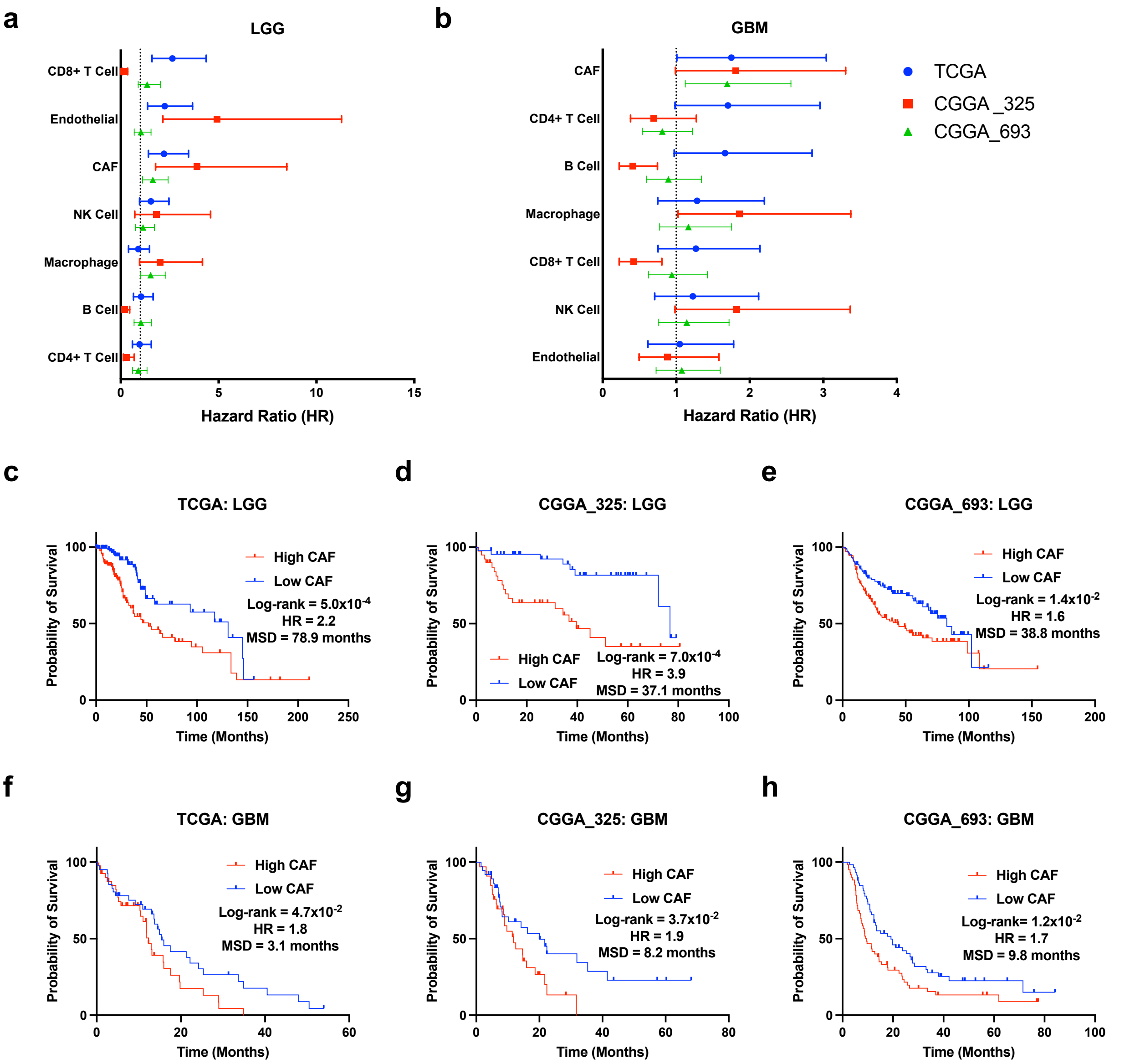
a
LGG
CD8+ T Cell
Endothelial
CAF
NK Cell
Macrophage
B Cell
CD4+ T Cell
0
5
10
15
Hazard Ratio (HR)
b
GBM
CAF
CD4+ T Cell
B Cell
Macrophage
CD8+ T Cell
NK Cell
Endothelial
0
1
2
3
4
Hazard Ratio (HR)
TCGA
CGGA_325
CGGA_693
c
TCGA: LGG
Probability of Survival
High CAF
Low CAF
Log-rank = 5.0x10-4
HR = 2.2
MSD = 78.9 months
Time (Months)
d
CGGA_325: LGG
High CAF
Low CAF
Log-rank = 7.0x10-4
HR = 3.9
MSD = 37.1 months
Time (Months)
e
CGGA_693: LGG
High CAF
Low CAF
Log-rank = 1.4x10-2
HR = 1.6
MSD = 38.8 months
Time (Months)
f
TCGA: GBM
High CAF
Low CAF
Log-rank = 4.7x10-2
HR = 1.8
MSD = 3.1 months
Time (Months)
g
CGGA_325: GBM
High CAF
Low CAF
Log-rank = 3.7x10-2
HR = 1.9
MSD = 8.2 months
Time (Months)
h
CGGA_693: GBM
High CAF
Low CAF
Log-rank= 1.2x10-2
HR = 1.7
MSD = 9.8 months
Time (Months)

Suppl Fig 1

**Supplementary S1. EPIC analysis of TCGA and CGGA glioma datasets.** (**a, b**) Hazard ratios associated with cell-type-specific estimated proportions across three independent data sets describing (**a**) LGG and (**b**) glioblastoma (data displayed as median with range). (**c-e**) Kaplan-Meier plots indicating percent survival over time (months) among LGG patients with either a high- or low-CAF abundance in the (**c**) TCGA (High CAF, n=127; Low CAF, n=127), (**d**) CGGA 325 (High CAF, n=45; Low CAF, n=45), and (**e**) CGGA 693 (High CAF, n=111; Low CAF, n=111) dataset (MSD, median survival difference; HR, hazard ratio). (**f-h**) Kaplan-Meier plots indicating percent survival over time (months) among glioblastoma patients with either a high- or low-CAF abundance in the (**f**) TCGA (High CAF, n=41; Low CAF, n=41), (**g**) CGGA 325 (High CAF, n=36; Low CAF, n=37), and (**h**) CGGA 693 datasets (High CAF, n=62; Low CAF, n=62).

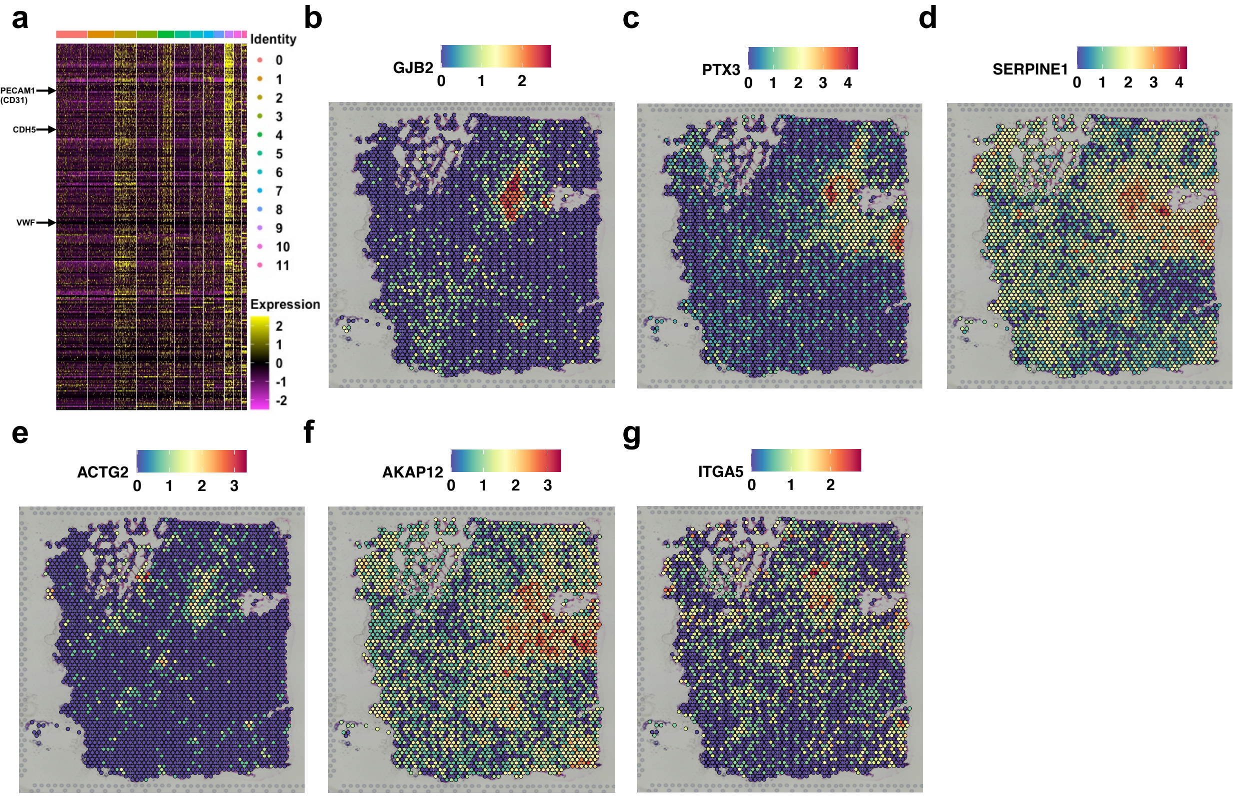

a
Identity
0
1
2
3
4
5
6
7
8
9
10
11
Expression
2
1
0
-1
-2
PECAM1 (CD31)
CDH5
VWF
b
GJB2
0 1 2
c
PTX3
0 1 2 3 4
d
SERPINE1
0 1 2 3 4
e
ACTG2
0 1 2 3
f
AKAP12
0 1 2 3
g
ITGA5
0 1 2


Suppl Fig 2

**Supplementary Figure S2, related to main figure 1. Expression patterns for selected genes or gene set in the spatial transcriptomics data.** (**a**) Heatmap indicating the expression of the signature genes previously associated with microvascular regions of proliferation are preferentially expressed in unique regions (2, 4, and 9) that are not enriched with CAFs. (**b-g**) Spatial dimensional plots showing gene expression patterns across human glioblastoma samples, (**b**) *GJB2*, (**c**) *PTX3*, (**d**) *SERPINE1*, (**e**) *ACTG2*, (**f**) *AKAP12*, and (**g**) *ITGA5*.

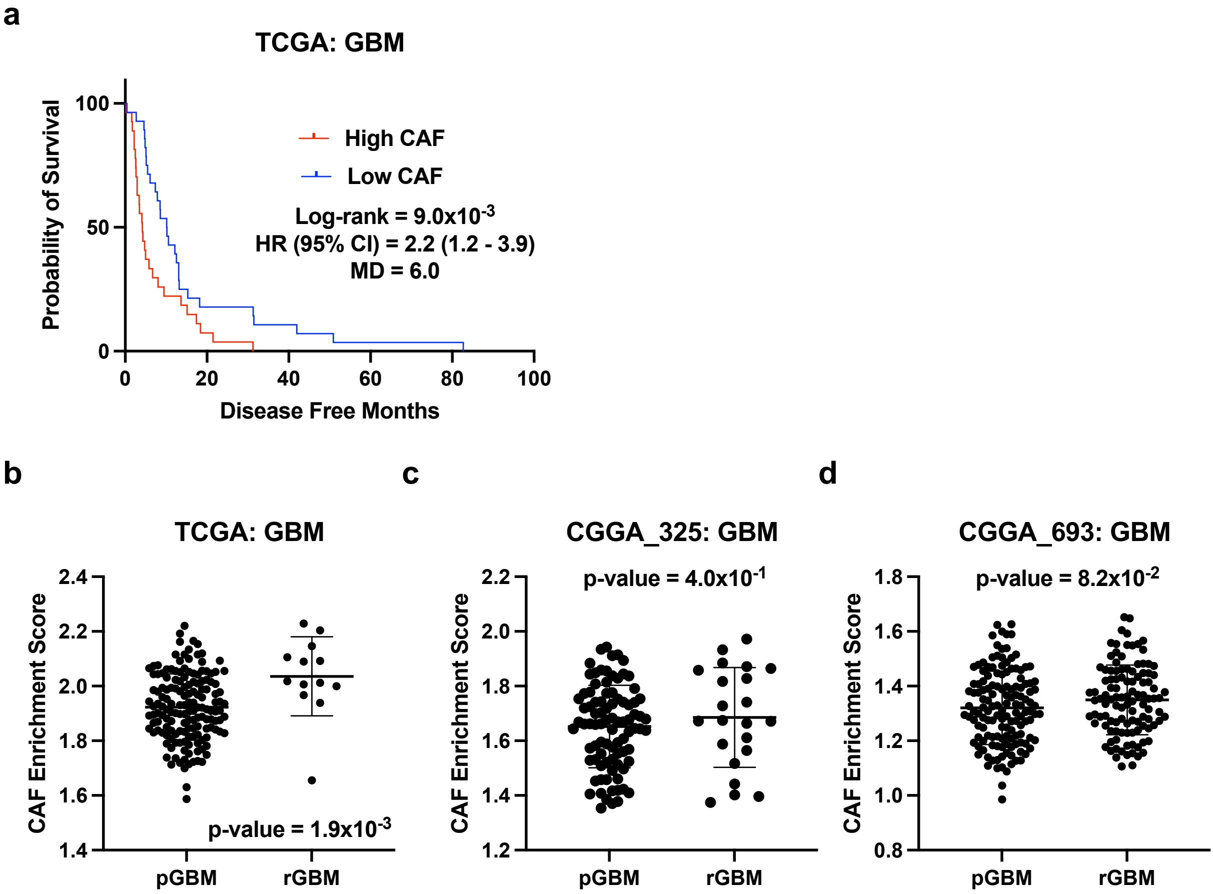
a
TCGA: GBM
High CAF
Low CAF
Log-rank = 9.0x10-3
HR (95% CI) = 2.2 (1.2 - 3.9)
MD = 6.0
Probability of Survival
Disease Free Months
b
TCGA: GBM
CAF Enrichment Score
p-value = 1.9x10-3
pGBM
rGBM
c
CGGA_325: GBM
p-value = 4.0x10-1
CAF Enrichment Score
pGBM
rGBM
d
CGGA_693: GBM
p-value = 8.2x10-2
CAF Enrichment Score
pGBM
rGBM

Suppl Fig 3

**Supplementary Figure S3, related to main figure 2. CAF enrichment score correlates with short disease-free progression.** (**a**) Kaplan-Meier plot indicating percent probability of event over time (disease free months) among glioblastomas patients in the TCGA cohort (High CAF, n=28; Low CAF, n=28). (**b-d**) Differences in CAF enrichment score between primary glioblastomas (pGBM) and reoccurring glioblastomas (rGBM) in (**b**) TCGA (pGBM, n=155; rGBM, n=13), (**c**) CGGA 325 (pGBM, n=88; rGBM, n=22), and (**d**) CGGA 693 (pGBM, n=140; rGBM, n=109) datasets (data displayed as mean ± S.D, p values from unpaired t test).

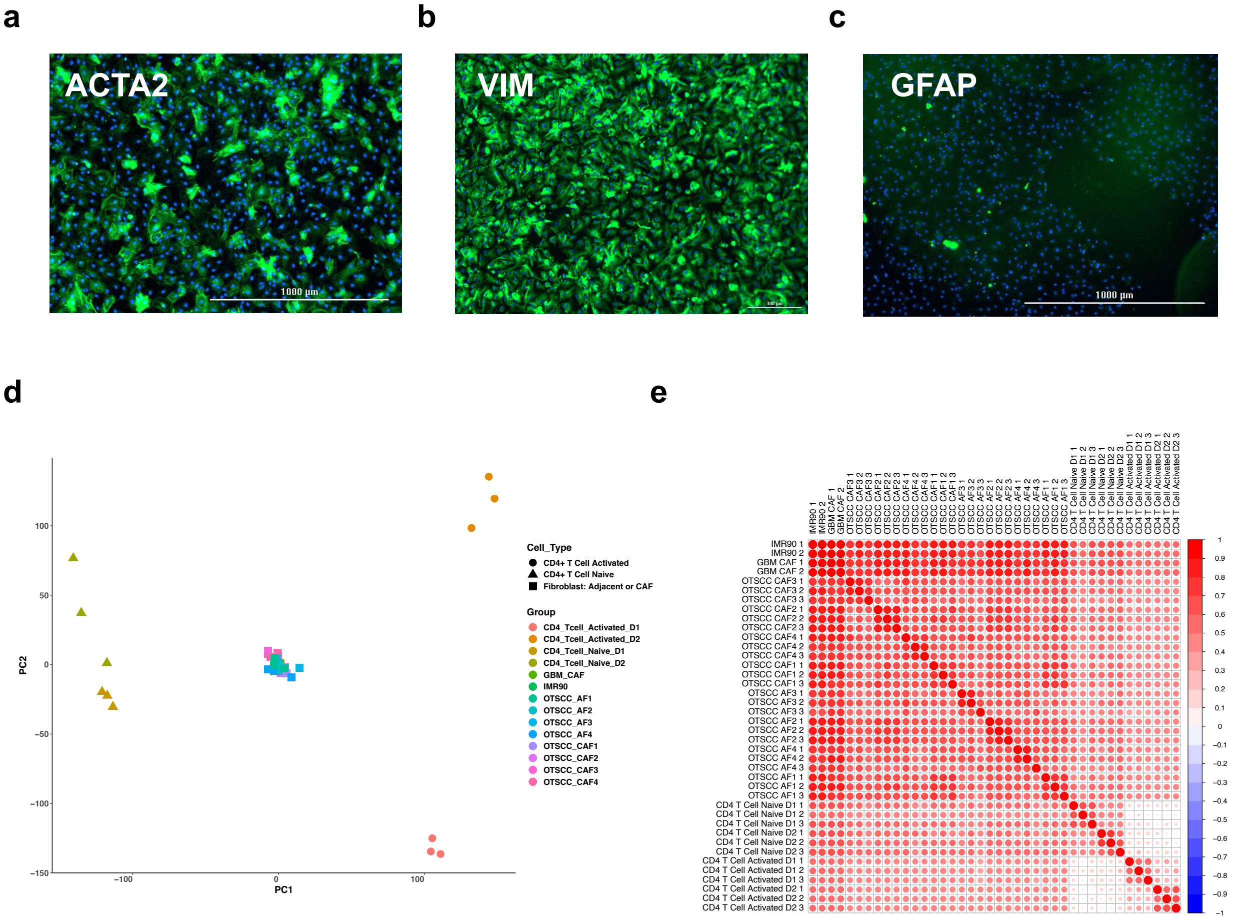
a
b
c
ACTA2
1000 µm
VIM
GFAP
1000 µm
d
e
PC2
PC1
100
50
0
−50
−100
−150
−100
0
100
Cell_Type
CD4+ T Cell Activated
CD4+ T Cell Naive
Fibroblast: Adjacent or CAF
Group
CD4_Tcell_Activated_D1
CD4_Tcell_Activated_D2
CD4_Tcell_Naive_D1
CD4_Tcell_Naive_D2
GBM_CAF
IMR90
OTSCC_AF1
OTSCC_AF2
OTSCC_AF3
OTSCC_AF4
OTSCC_CAF1
OTSCC_CAF2
OTSCC_CAF3
OTSCC_CAF4
IMR90 1
IMR90 2
GBM CAF 1
GBM CAF 2
OTSCC CAF3 1
OTSCC CAF3 2
OTSCC CAF3 3
OTSCC CAF2 1
OTSCC CAF2 2
OTSCC CAF2 3
OTSCC CAF4 1
OTSCC CAF4 2
OTSCC CAF4 3
OTSCC CAF1 1
OTSCC CAF1 2
OTSCC CAF1 3
OTSCC AF3 1
OTSCC AF3 2
OTSCC AF3 3
OTSCC AF2 1
OTSCC AF2 2
OTSCC AF2 3
OTSCC AF4 1
OTSCC AF4 2
OTSCC AF4 3
OTSCC AF1 1
OTSCC AF1 2
OTSCC AF1 3
CD4 T Cell Naive D1 1
CD4 T Cell Naive D1 2
CD4 T Cell Naive D1 3
CD4 T Cell Naive D2 1
CD4 T Cell Naive D2 2
CD4 T Cell Naive D2 3
CD4 T Cell Activated D1 1
CD4 T Cell Activated D1 2
CD4 T Cell Activated D1 3
CD4 T Cell Activated D2 1
CD4 T Cell Activated D2 2
CD4 T Cell Activated D2 3
1
0.9
0.8
0.7
0.6
0.5
0.4
0.3
0.2
0.1
0
−0.1
−0.2
−0.3
−0.4
−0.5
−0.6
−0.7
−0.8
−0.9
−1

### Suppl Fig 4

**Supplementary Figure S4, related to main figure 3. Validation of glioblastoma-derived CAF cell line. (a-c)** Immunocytochemistry staining on the GBM-CAF cell line showing the expression of (**a**) ACTA2, (**b**) VIM, and (**c**) GFAP. Scale bars, 1000 µm. (**d**) PCA of proteomic data generated from whole cell lysates of glioblastoma-derived CAF cell lines (n=2), IMR90 (n=2), CAFs (n=12) and adjacent fibroblasts (n=12) isolated from human OTSCC tumors, and naïve (n=3) and activated (n=3) CD4+ T cells isolated from two separate donors. (**e**) Heatmap depicting correlation among proteomic datasets in D.

a

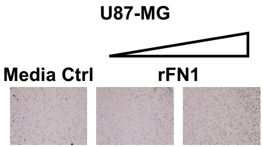


b

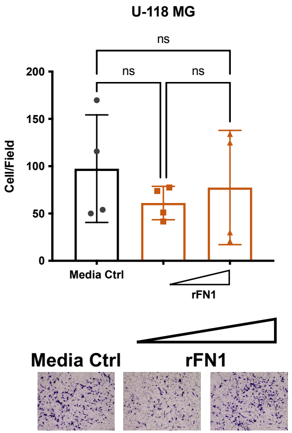


c

| | Fibro$^{CM}$ | | CAF$^{CM}$ | | rFN1 | | |
|---|---|---|---|---|---|---|---|
| | Migration | Invasion | Migration | Invasion | Migration | Invasion | **Final Status:** |
| T98-G | ✅ | ✅ | ✅ | ✅ | ✅ | ✅ | Responder |
| LN-229 | ✅ | ✅ | ✅ | ✅ | ✅ | ✅ | Responder |
| U87-MG | ✅ | ❌ | ✅ | ❌ | ✅ | ❌ | Partial-Responder |
| U-118 MG | ❌ | ❌ | ❌ | ❌ | ✅ | ❌ | Partial-Responder |

**Suppl Fig 5**

**Figure S5, related to main figure 4. Fibro$^{CM}$, CAF$^{CM}$, and FN1 showed variable effects on glioblastoma cell lines. (a-b)** Transwell invasion results for glioblastoma cell lines exposed to human recombinant FN1, (**a**) U87-MG and (**b**) U-118 MG. Data were generated from n=3 independent experiments and displayed as mean ± S.D. (**c**) A table summarizing the effects of Fibro$^{CM}$, CAF$^{CM}$, or FN1 on the migration and invasion of four glioblastoma cell lines.

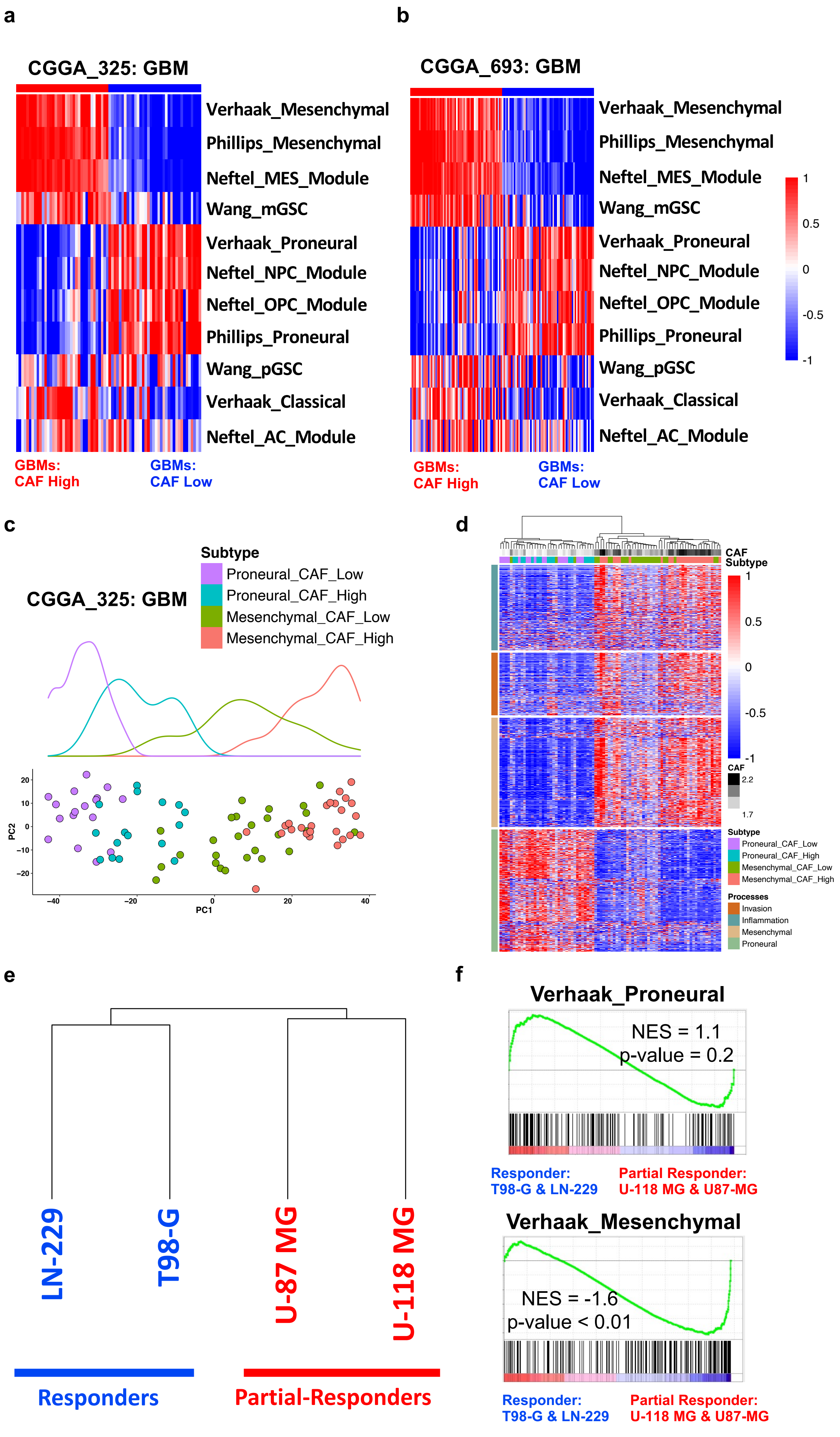

a
CGGA_325: GBM
Verhaak_Mesenchymal
Phillips_Mesenchymal
Neftel_MES_Module
Wang_mGSC
Verhaak_Proneural
Neftel_NPC_Module
Neftel_OPC_Module
Phillips_Proneural
Wang_pGSC
Verhaak_Classical
Neftel_AC_Module
GBMs: CAF High
GBMs: CAF Low
b
CGGA_693: GBM
Verhaak_Mesenchymal
Phillips_Mesenchymal
Neftel_MES_Module
Wang_mGSC
Verhaak_Proneural
Neftel_NPC_Module
Neftel_OPC_Module
Phillips_Proneural
Wang_pGSC
Verhaak_Classical
Neftel_AC_Module
1
0.5
0
-0.5
-1
GBMs: CAF High
GBMs: CAF Low
c
CGGA_325: GBM
Subtype
Proneural_CAF_Low
Proneural_CAF_High
Mesenchymal_CAF_Low
Mesenchymal_CAF_High
PC2
PC1
20
10
0
−10
−20
−40
−20
0
20
40
d
CAF
Subtype
1
0.5
0
-0.5
-1
CAF
2.2
1.7
Subtype
Proneural_CAF_Low
Proneural_CAF_High
Mesenchymal_CAF_Low
Mesenchymal_CAF_High
Processes
Invasion
Inflammation
Mesenchymal
Proneural
e
LN-229
T98-G
U-87 MG
U-118 MG
Responders
Partial-Responders
f
Verhaak_Proneural
NES = 1.1
p-value = 0.2
Responder: T98-G & LN-229
Partial Responder: U-118 MG & U87-MG
Verhaak_Mesenchymal
NES = -1.6
p-value < 0.01
Responder: T98-G & LN-229
Partial Responder: U-118 MG & U87-MG


**Suppl Fig 6**

**Supplementary S6, related to main figure 5. Analysis of the CGGA cohort supports the hypothesis that CAFs facilitate a proneural-to-mesenchymal transition in glioblastoma.** Heatmap depicting individual enrichment scores for signature gene sets describing specific glioblastoma subtypes among glioblastoma samples (columns) with a high and low CAF enrichment score in the (**a**) CGGA 325 and (**b**) CGGA 693 datasets. (**c**) PCA plot for glioblastoma samples from the CGGA 325 cohort, with colors for four groups: proneural CAF low (n=17), proneural CAF high (n=16), mesenchymal CAF low (n=26), and mesenchymal CAF high (n=25). The densities of samples are shown above. (**d**) Heatmap showing changes in the expression of genes associated with inflammation, invasion, mesenchymal-subtype, and proneural-subtype as glioblastomas accumulate more CAFs in the CGGA 325 cohort. (**e**) Dendrogram showing gene expression correlation of four glioma cell-lines. (**f**) GSEA enrichment plots for the proneural and mesenchymal glioblastoma subtype gene sets, comparing responder cell lines (T98-G and LN-229) and partial responder cell lines (U118-MG and U87-MG) in the RNA-seq data.

**Suppl Table 1. Multivariate survival analysis models and association of CAFs with clinical and molecular features of both LGG and GBM patients.**

| | | TCGA: LGG and GBM (n=672) | | CGGA_325: LGG and GBM (n=325) | | CGGA_693: LGG and GBM (n=693) | |
|---|---|---|---|---|---|---|---|
| **Predictor** | **Levels** | **HR (95% CI)** | **p-value** | **HR (95% CI)** | **p-value** | **HR (95% CI)** | **p-value** |
| Age | Continuous | 1.03 (1.01 - 1.05) | 0.001 | 1.0 (0.9 - 1.02) | 0.8 | 1.0 (0.9 - 1.01) | 0.3 |
| Gender | Female | 1 (ref) | | 1 (ref) | | 1 (ref) | |
| | Male | 1.0 (0.7 - 1.4) | 0.9 | 1.05 (0.9 - 1.02) | 0.8 | 1.02 (0.8 - 1.3) | 0.9 |
| IDH Mutation | WT | 1 (ref) | | 1 (ref) | | 1 (ref) | |
| | Mutant | 0.3 (0.1 - 0.8) | 0.01 | 0.7 (0.4 - 1.1) | 0.2 | 0.5 (0.4 - 0.7) | < 0.001 |
| Neoplasm Grade | 2 | 1 (ref) | | 1 (ref) | | 1 (ref) | |
| | 3 | 1.2 (0.4 - 3.3) | 0.8 | 4.6 (2.5 - 8.6) | < 0.001 | 2.8 (2.0 - 3.9) | < 0.001 |
| | 4 | 2.1 (0.7 - 7.0) | 0.2 | 9.0 (4.7 - 17.0) | < 0.001 | 4.5 (3.1 - 6.6) | < 0.001 |
| CAF Enrichment Score | Continuous | 6.4 (1.1 - 36.2) | 0.04 | 5.4 (1.5 - 19.2) | 0.01 | 3.3 (1.3 - 8.1) | 0.01 |

**Suppl Table 2. Multivariate survival analysis models and association of CAFs with clinical and molecular features of LGG patients.**

| | | TCGA: LGG (n=511) | | CGGA_325: LGG (n=181) | | CGGA_693: LGG (n=433) | |
|---|---|---|---|---|---|---|---|
| **Predictor** | **Levels** | **HR (95% CI)** | **p-value** | **HR (95% CI)** | **p-value** | **HR (95% CI)** | **p-value** |
| Age | Continuous | 1.1 (1.0 - 10.2) | 0.02 | 1.0 (0.9 - 1.0) | 0.4 | 1.0 (0.9 - 1.0) | 0.7 |
| Gender | Female | 1 (ref) | | 1 (ref) | | 1 (ref) | |
| | Male | 1.2 (0.4 - 3.3) | 0.7 | 0.8 (0.5 - 1.6) | 0.6 | 1.2 (0.9 - 1.6) | 0.3 |
| IDH Mutation | Mutant | 1 (ref) | | 1 (ref) | | 1 (ref) | |
| | WT | 3.2 (1.0 - 10.2) | 0.05 | 2.2 (1.1 - 4.4) | 0.02 | 2.0 (1.4 - 2.8) | <0.0001 |
| Neoplasm Grade | 2 | 1 (ref) | | 1 (ref) | | 1 (ref) | |
| | 3 | 0.9 (0.3 - 2.9) | 0.8 | 3.8 (1.8 - 7.9) | <0.0001 | 3.0 ( 2.1 - 4.2) | <0.0001 |
| CAF Enrichment Score | Continuous | 1339.0 (10.8 - 166054.1) | 0.003 | 10.2 (1.0 - 104.0) | 0.049 | 12.4 (3.2 - 48.0 | <0.0001 |

**Suppl Table 3. Multivariate survival analysis models and association of CAFs with clinical and molecular features of GBM patients.**

| | | TCGA: GBM (n=161) | | CGGA_325: GBM (n=144) | | CGGA_693: GBM (n=249) | |
|---|---|---|---|---|---|---|---|
| **Predictor** | **Levels** | **HR (95% CI)** | **p-value** | **HR (95% CI)** | **p-value** | **HR (95% CI)** | **p-value** |
| Age | Continuous | 1.05 (1.02 - 1.0) | < 0.0001 | 1.0 (0.9 -1.0) | 0.7 | 1.0 (0.9-1.0) | 0.2 |
| Gender | Female | 1 (ref) | | 1 (ref) | | 1 (ref) | |
| | Male | 1.0 (0.6 - 1.7) | 0.9 | 1.2 (0.8 - 2.0) | 0.4 | 1.1 (0.8 - 1.4) | 0.6 |
| IDH Mutation | Mutant | 1 (ref) | | 1 (ref) | | 1 (ref) | |
| | WT | 1.9 (0.2 - 17.5) | 0.6 | 1.1 (0.6 - 2.0) | 0.4 | 1.6 (1.0 - 2.4) | 0.03 |
| MGMT Promoter | Unmethylated | 1 (ref) | | NA | NA | NA | NA |
| | Methylated | 0.8 (0.5 - 1.3) | 0.3 | NA | NA | NA | NA |
| CAF Enrichment Score | Continuous | 7.5 (0.8 - 69.3) | 0.08 | 5.8 ( 1.2 - 29.0) | 0.03 | 1.6 (0.5 - 5.2) | 0.4 |
